# Metabolomic, photoprotective, and photosynthetic acclimatory responses to post-flowering drought in sorghum

**DOI:** 10.1101/2022.01.14.476420

**Authors:** Christopher R. Baker, Dhruv Patel, Benjamin J. Cole, Lindsey G. Ching, Oliver Dautermann, Armen C. Kelikian, Cayci Allison, Julie Pedraza, Julie Sievert, Aivett Bilbao, Joon-Yong Lee, Young-Mo Kim, Jennifer E. Kyle, Kent J. Bloodsworth, Vanessa Paurus, Kim K. Hixson, Robert Hutmacher, Jeffery Dahlberg, Peggy G. Lemaux, Krishna K. Niyogi

## Abstract

Climate change is globally affecting rainfall patterns, necessitating the improvement of drought tolerance in crops. *Sorghum bicolor* is a drought-tolerant cereal capable of producing high yields under water scarcity conditions. Functional stay-green sorghum genotypes can maintain green leaf area and efficient grain filling in terminal post-flowering water deprivation, a period of ~10 weeks. To obtain molecular insights into these characteristics, two drought-tolerant genotypes, BTx642 and RTx430, were grown in control and terminal post-flowering drought field plots in the Central Valley of California. Photosynthetic, photoprotective, water dynamics, and biomass traits were quantified and correlated with metabolomic data collected from leaves, stems, and roots at multiple timepoints during drought. Physiological and metabolomic data was then compared to longitudinal RNA sequencing data collected from these two genotypes. The metabolic response to drought highlights the uniqueness of the post-flowering drought acclimation relative to pre-flowering drought. The functional stay-green genotype BTx642 specifically induced photoprotective responses in post-flowering drought supporting a putative role for photoprotection in the molecular basis of the functional stay-green trait. Specific genes are highlighted that may contribute to post-flowering drought tolerance and that can be targeted in crops to maximize yields under limited water input conditions.

**Highlight:** Pathways contributing to the long-term maintenance of photosynthetic activity in terminal post-flowering drought are revealed by a comprehensive approach combining in-field photosynthetic physiological analysis, metabolomics, and transcriptomics.

## Introduction

Worldwide, drought remains the primary abiotic cause of agricultural yield loss, and climate change may accelerate the impact of drought on agriculture as the frequency and severity of droughts increase (Lesk *et al*., 2016). The overuse of groundwater, largely driven by agricultural demand (Giordano, 2009; Giordano *et al*., 2019), also limits irrigation as a long-term solution to maintaining agricultural productivity in a world experiencing hotter temperatures (Lobell *et al*., 2014; Ort and Long, 2014). Defining and tweaking the molecular mechanisms underlying drought-adapted traits in plants is vital to maintaining high yields under expected future climatic conditions (Varshney *et al*., 2018).

Drought tolerance is a complex, quantitative trait dependent on plant developmental stage and the severity of the water deficit (Luo *et al*., 2019). Crops, like sorghum, that perform C4 photosynthesis, an evolutionary innovation in the carbon (C) reactions of photosynthesis and anatomy of the leaf tissue that increases intrinsic water use-efficiency (WUE_i_) by reducing transpirational loss, can exhibit higher drought tolerance than those using C3 photosynthesis. Of the C4 crops, sorghum *[Sorghum bicolor* (L.) Moench] is exceptionally drought-tolerant (Kimber, 2000), and the timing of drought before anthesis (pre-flowering drought) or post-anthesis (post-flowering drought) has markedly different outcomes (Rosenow and Clark 1995; Rosenow et al. 1996; Varoquaux et al. 2019). In the case of post-flowering drought stress, stalk-lodging rates and leaf senescence can increase and, of agronomic importance, grain size and grain yield can be decreased (Thomas and Howarth, 2000).

Notably, the extent of post-flowering drought tolerance also differs between sorghum genotypes with so-called “stay-green” genotypes able to delay the senescence of the upper canopy until after the final stages of grain filling (Krieg and Hutmacher, 1986; Borrell *et al*., 2000). In “functional stay-green” plants, such as the sorghum genotype BTx642, delayed leaf senescence in terminal post-anthesis water deprivation is part of a suite of advantageous traits contributing to maintenance of high grain yields and grain size and prevention of stalk lodging (Tuinstra *et al*., 1997; Thomas and Howarth, 2000; Harris *et al*., 2007). This is contrasted with so-called “cosmetic stay-green” plants, which block chlorophyll degradation and, thus, remain green in drought but do not maintain high yields (Thomas and Howarth, 2000; Hörtensteiner and Kräutler, 2011).

At the whole-plant level, BTx642 has low tillering rates and less above-ground biomass per plant relative to post-flowering drought-susceptible sorghum genotypes at anthesis (Borrell *et al*., 2014*b*,*a*). At the cellular level, stay-green sorghum genotypes maintain photosynthetic machinery through the grain-filling period in post-flowering drought (Borrell *et al*., 2001; Varoquaux *et al*., 2019), which may contribute to efficient grain filling and to preventing stalk lodging by maintaining high sugar levels in the stalk (Rosenow and Clark, 1995; Sanchez *et al*., 2002). Further, maintenance of photosynthetic leaf area during post-flowering drought will only be beneficial to the genotype if sufficient water reserves are available to allow stomata to remain partly open for CO_2_ assimilation (Borrell *et al*., 2001; Kamal *et al*., 2019; Varoquaux *et al*., 2019).

Leaf senescence is a developmentally controlled response that is responsive to abiotic stress signals. In particular, elevated reactive oxygen species (ROS) levels can induce early leaf senescence in drought (Cruz de Carvalho, 2008; Noctor *et al*., 2014). Excess excitation energy in drought drives ROS production leading to the peroxidation of polyunsaturated lipids, damage to proteins, and the inactivation of pigments and antioxidants. Plants have evolved a suite of photoprotective responses to manage ROS (Li *et al*., 2009). These include photoprotective antioxidants in photosynthetic and epidermal tissues, such as ascorbate, tocopherols, and photoprotective flavonoids (Logan *et al*., 2006; Li *et al*., 2009; Agati and Tattini, 2010), as well as activation of non-photochemical quenching (NPQ), the controlled dissipation of excess excitation energy as heat (Cousins *et al*., 2002; Golding and Johnson, 2003; Jung, 2004; Ogbaga *et al*., 2014; Lima Neto *et al*., 2017). Thus, strong photoprotective responses may act as a key post-flowering drought tolerance trait, however, direct evidence is lacking for this hypothesis.

The molecular response of sorghum to post-flowering drought in the field has not been extensively characterized. In a recent study, the time-resolved transcriptomic response was determined for pre-flowering and post-flowering droughted field-grown sorghum genotypes BTx642 and RTx430 (Varoquaux *et al*., 2019). Paralleling this study, two drought-tolerant sorghum genotypes, BTx642 and RTx430, were grown in irrigated plots in the California Central Valley in 2019 under both control and post-flowering drought conditions. BTx642 was selected as a functional stay-green variety, whereas RTx430 has strong drought tolerance but lacks the full suite of stay-green traits (Crasta et al. 1999). The aim of this study was to: (1) identify shared changes in metabolite and lipid levels in leaves, stems, and roots in response to post-flowering drought, (2) measure the extent of drought-induced stomatal closure in the two genotypes and identify stomatal closure regulators linked to this response, and (3) determine whether post-flowering drought-induced photoprotective responses, particularly in the stay-green genotype BTx642, contribute to the maintenance of green leaf area in a stay-green genotype under field conditions. To this end, photosynthetic, photoprotective, and water dynamics traits under the two field growth conditions were quantified in both genotypes across multiple drought timepoints and samples were harvested for metabolomic and lipidomic analysis. Physiological and metabolomic datasets were then compared to transcriptomic data collected from the same genotypes under identical growth conditions in a prior year (Varoquaux *et al*., 2019) in order to identify specific candidate genes that may contribute to post-flowering drought tolerance in sorghum. In particular, candidate genes are identified that may contribute to the stronger photoprotective responses and the control of stomatal closure during drought, as discovered in this work.

## Materials and Methods

### Field growth conditions

Sorghum genotypes BTx642 and RTx430 were grown in Parlier, CA (36.6008°N, 119.5109°W) in 2019 in a Hanford sandy loam soil (pH = 7.37) with a silky substratum in 0.071 hectares (ha) plots of ten rows each. Two watering conditions were used on plots: I) control, consisting of weekly watering five days prior to sampling dates, with the first irrigation starting 18 days after planting (DAP) and continuing until 123 DAP and II) post-flowering drought, consisting of regular irrigation up through and including irrigation before 65 DAP– at which point over 50% of the plants flowered – with terminal water deprivation from that point onwards (Figure 1B). Pre-planting irrigation was performed for all plots such that the upper 122 cm of soil would have been refilled to soil field capacity. Plots receiving water were irrigated at seven-day intervals using drip irrigation lines placed on the soil surface of each furrow. All irrigated plots received equal volumes of water equal to 100% of the average weekly calculated crop evapotranspiration for the 7-day period before irrigation. Irrigation once per week in plots replicates sorghum farming irrigation practices in the Western United States. Additionally, providing equal water volume to all irrigated plots prevents a scenario where genotypic differences in evapotranspiration rates lead to a difference in total water volume supplied to specific plots. Total final biomass was comparable between control plots for both genotypes with an average forage (65% moisture) of 13.02 T ac^−1^ for BTx642 and 13.28 T ac^−1^ for RTx430. For greater details on crop evapotranspiration and irrigation management, see supplemental materials and Xu et al. (Xu *et al*., 2018).

**Figure 1:**
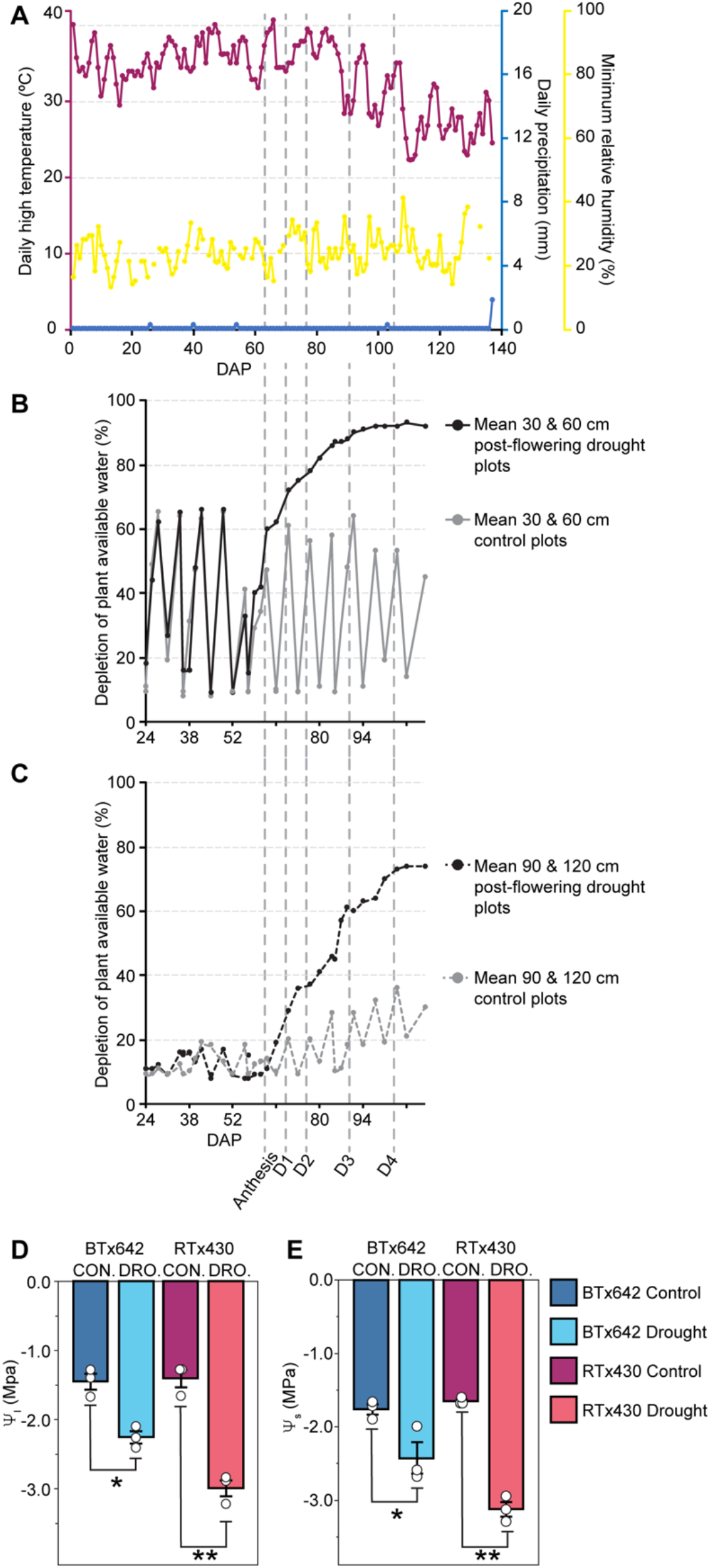
Field conditions, soil water depletion, and leaf water potential response to terminal drought stress. (A) Data collected from June 11 to October 26, 2019 at Parlier Weather Station A (Parlier, CA, USA). Daily high temperature (axis 1, magenta), daily precipitation (axis 2, blue), and minimum relative humidity (axis 3, yellow). The y-axis upper bound for each variable is set to the daily annual maximum value for 2019. (B-C) Soil water data expressed as percent plant available soil water depletion between −0.02 MPa (field capacity) and −1.5 MPa (permanent wilting point). Drought plots (dark grey) and control plots (light grey) with sensors at (B) 30 and 60 cm (solid lines) and (C) 90 and 120 cm depth (dashed lines). Sampling dates are labeled as D1 through D4. (D) Midday leaf water potential (Ψ_I_) and (E) osmotic potential (Ψ_S_) collected on D4 (40 days without water). BTx642 control (dark blue), BTx642 drought (light blue), RTx430 control (purple), and RTx430 drought (pink). Mean values ± standard errors (*n* = 3 plots) with mean values for each individual plot displayed as dots (white). Significant differences as measured by a two-tailed *t*-test for control vs. treatment pairs are indicated by asterisks (* < 0.05, ** < 0.005).

Planting in 2019 occurred on June 10. Four sampling dates were selected and for each date, control plots had not received water for 5 days: 1) August 20, 2019 (D1), five days since last watering for all plots, 70 DAP, II) August 27, 2019 (D2), 12 days of post-flowering drought, 77 DAP, 3) September 10, 2019 (D3), 26 days of post-flowering drought, 91 DAP, 4) September 24, 2019 (D4), 40 days of post-flowering drought plots, 105 DAP.

### Harvest data

Harvest data metrics of 50% flowering time, plant height, 1000 seed weight, seed weight (13% moisture) ha^−1^, and forage (65% moisture) weight ha^−1^ were determined for each plot as previously described (Xu *et al*., 2018). Three replicate plots were planted for each genotype under the same treatment regimes.

### Leaf phenotypic traits

On each of the four 2019 sampling dates, gas exchange and chlorophyll fluorescence measurements were collected within two time windows: 9:30 to 11:00 (morning) and 14:00 to 16:00 (mid-afternoon) using LI-COR 6400XT instruments (LI-COR, Lincoln, NE, USA). Given that these sampling dates all occurred post-anthesis, all leaves had emerged and thus, it was possible to randomly sample the uppermost three leaves including the flag leaf from plants growing in the interior of each plot at each sampling date. Each of the LI-COR 6400XT instruments were factory calibrated the month prior to this field work, and the calibrations and instrument checks as described in Chapter 4 of the LI-COR 6400 manual were performed on each sampling date. Leaves were maintained near ambient light levels and temperatures by measuring ambient PAR levels and local temperatures and re-adjusting actinic light levels and blocking temperature prior to each set of measurements. The ratio of blue-to-red LED contribution to the cuvette light source was 10% / 90%. Relative humidity in the measurement cuvette was maintained between 50% to 60% to maintain stomatal aperture width. Flow rate was set to 400 μmol s^−1^ and sample [CO_2_] to 400 μmol mol^−1^. Stability variables typically converged within 60 s of clamping a leaf, then an infrared gas analyzer match was performed and, once stability variables were restored following the match, the measurement was taken. Leaves were clamped to avoid the midrib and always near the midpoint of the leaf (*i.e*., equal distance from the tip and leaf base). A multiphase flash routine was used to estimate chlorophyll fluorescence parameters (Loriaux *et al*., 2013). Prior to the measurement of F_o_’, fluorescence level of a light acclimated sample when all PSII reaction centers are open, a far-red light pulse was given to leaves of 25 μmol photons m^−2^ s^−1^ for 1 s prior to actinic light switching and then lasting an additional 5 s and ending 1 s prior to the measurement. A minimum of eight leaves were randomly sampled per plot per timepoint.

Leaf water potential (ψ_I_) and osmotic potential (ψ_s_) were measured on the uppermost non-flag leaf of the main culm from three randomly selected plants from the interior of each plot in mid-afternoon the day after D4 (on September 24, 2019, 105 DAP). Thus, the control plots had not received water for six days and post-flowering droughted plots had not received water for 41 days. On this same day (105 DAP), green leaf area images were collected and F_v_/F_m_ and NPQ were determined. Specific to NPQ measurements, these values were measured exclusively on leaves without visible signs of leaf senescence in both control and droughted plots. This decision was made to ensure that photoprotective traits could be accurately quantified in leaves with photosynthetic machinery intact prior to the onset of leaf senescence traits. Green leaf area was determined by imaging the three uppermost leaves including the flag leaf on ten randomly selected plants per plot. Stomatal density and guard cell length were quantified using leaf peels collected on the D4 sampling date from the abaxial leaf surface of the uppermost non-flag leaf of the main culm from (Lopez *et al*., 2017). More details of leaf phenotypic measurements can be found in the supplemental materials.

### Sample collection and processing

Within each plot, samples of leaves, stems, and roots from individual plants were manually collected on the same day of the week and time of the day for each sampling date. Three plants from each plot were collected and the uppermost three leaves, stems (below the peduncle and above the node for the next leaf below), and roots were harvested to create a single leaf, stem, and root sample for each plot for each timepoint. Individual sorghum plants were harvested at various developmental stages using a shovel to a depth of approximately 30 cm. All samples were then flash-frozen in liquid nitrogen within 5 minutes of being removed from the field. Root tissue was collected as previously described (Xu *et al*., 2018). Each week, all samples were collected less than 1 hr after dawn (dawn), within 1 hr of the midpoint of the light period (midday), and less than 1 hr before dusk (dusk).

### Metabolite extraction, quantification, and metabolomics

For details of metabolite extractions and spectrophotometric quantification of specific metabolites see supplemental materials. Leaf tissue samples from sampling dates D2, D3, and D4 were analyzed by GC-MS, lipidomics, and SPE-IMS-MS. Metabolomic data were collected for stem and root samples from D2, D3, and D4 sampling dates exclusively by IMS. For GC-MS, MPLEx extraction was applied to the samples which were weighed at 1 g (Nakayasu *et al*., 2016). Then, samples were completely dried under a speed vacuum concentrator. Dried metabolites were chemically derivatized and analyzed as reported previously (Kim *et al*., 2015) and further described in supplemental materials. Metabolites were initially identified by matching experimental spectra to an augmented version of the Agilent Fiehn Metabolomics Library, containing spectra and validated retention indices for almost 1000 metabolites (Kind *et al*., 2009) and additionally cross-checked by matching with NIST17 GC/MS Spectral Library and Wiley Registry 11th edition. All metabolite identifications were manually validated to minimize deconvolution and identification errors during the automated data processing. Data were log_2_ transformed and then mean-centered across the log_2_ distribution. C and N values were determined at the Center for Stable Isotope Biogeochemistry at UC-Berkeley using leaf samples from the D4 time point. Organic nitrogen (*N_org_*) values were calculated by subtracting total N levels by spectrophotometrically determined ammonium (Ammonia assay kit, Megazyme, Bray, Ireland) and nitrate levels (Bloom *et al*., 2014).

For lipidomics, total lipid extracts (TLEs) were analyzed as outlined in Kyle et al. (2017) and further detailed in the supplemental materials. TLEs were analyzed in both positive and negative electrospray ionization modes, and lipids were fragmented using alternating higher-energy collision dissociation (HCD) and collision-induced dissociation (CID) (Kyle *et al*., 2017). Identifications were made using LIQUID (Kyle *et al*., 2017) and manually validated by examining the MS/MS spectra for fragment ions characteristic of the classes and acyl chain compositions of the identified lipids. In addition, the precursor ion isotopic profile extracted ion chromatogram, and mass measurement error along with the elution time were evaluated. All LC-MS/MS data were aligned and gap-filled to this target library for feature identification using MZmine 2 (Pluskal *et al*., 2010), based on the identified lipid name, observed m/z, and retention time. Data from each ionization mode were aligned and gap-filled separately. Aligned features were manually verified and peak apex intensity values were exported for statistical analysis.

For SPE-IMS-MS Metabolomics, extracts were analyzed using a RapidFire 365 (Zhang *et al*., 2016) coupled with an Agilent 6560 Ion Mobility QTOF MS system (Agilent Technologies, Santa Clara, CA, USA) and described in detail in the supplemental materials. The PNNL-PreProcessor v2020.07.24 (https://omics.pnl.gov/software/pnnl-preprocessor) was used to generate new raw MS files (Agilent MassHunter “.d”) for each sample, run with all frames (ion mobility separations) summed into a single frame and applying 3-points smoothing in the ion mobility dimension and noise filtering with a minimum intensity threshold of 20 counts. Details of the data processing and compound identification can be found in supplemental materials.

### Statistical analysis

All statistical analyses, excepting metabolomics and lipidomics data, were performed using JMP Pro 16 software (JMP, Cary, NC, USA). Prior to the analysis of gas exchange values, six measurements (out of the 462 measurements taken) with physiologically impossible *C_i_* values were removed from our datasets and attributed to either machine or user error. Metaboanalyst 5.0 (Chong *et al*., 2018) was used to guide the selection of metabolites to further analyze using a significance threshold of *p* < 0.01 for the significance of the differential abundance of drought vs. control on at least one sampling date. Ontology enrichment for differentially abundant lipids across comparisons was performed using Lipid Mini-On (Clair *et al*., 2019).

### Transcriptomic data processing and visualization

To generate expression plots for selected gene sets, we obtained normalized counts of *S. bicolor* genes mapped to a common reference (*S. bicolor* BTx623), and accompanying metadata from the EPICON field trial described previously (Varoquaux *et al*., 2019). Normalized counts were then summarized for control-treated leaf samples for each genotype, week, and gene by taking the arithmetic mean (*n* = 1-3), and Log_2_-transformation (with a pseudocount of 1). These values were subtracted from Log_2_-transformed (plus a pseudocount of 1) normalized counts for each locus, genotype, day, and treatment from the EPICON dataset, to generate a control mean-corrected dataset of gene expression for pre- and post-flowering drought treatments. These values were then plotted as points, with loess-smoothed values computed from these transformed data plotted as lines.

## Combined results and discussion

### Induction of post-flowering drought response in the field

BTx642 and RTx430 plants reached 50% inflorescence emergence by 69 days after planting (DAP) and 71 DAP, respectively (Fig. 1, Table 1; see Fig. S1 for details of the field layout). Before anthesis, the average maximum daily temperature was 35.5°C with a range from 29.4°C to 40.0°C for maximum daily temperatures (Fig. 1A). Post-anthesis temperatures declined moderately with an average maximum daily temperature of 34.6°C with a range of 26.7°C to 40.6°C for maximum daily temperatures throughout the grain-filling period. Relative humidity was in general low with an average minimum daily value of 23.9% with a range of 13% to 35% from the time of germination to the end of the grain filling period (Fig. 1A). No precipitation occurred during the growth lifecycle (Fig. 1A).

**Table 1:** Harvest and selected leaf phenotypic responses to terminal post-flowering drought

| Variable | BTx642 |  |  | RTx430 |  |  | One-way ANOVA |  |
| --- | --- | --- | --- | --- | --- | --- | --- | --- |
| | Control | Drought | $\Delta$<br>(Drought - Control) | Control | Drought | $\Delta$<br>(Drought - Control) | BTx642<br>(Drought vs. control) | RTx430<br>(Drought vs. control) |
| Post-flowering drought tolerance traits |  |  |  |  |  |  |  |  |
| Flowering time (DAP) | 69.33 | 67.00 | -2.33 | 70.67 | 70.67 | 0 | n.s. | n.s. |
| Forage (65% moisture) (Mg ha<sup>-1</sup>) | 29.19 | 26.32 | -2.87 | 29.77 | 24.01 | -5.76 | n.s. | n.s. |
| 1000 seed (g) | 26.70 | 28.87 | 2.17 | 30.55 | 27.98 | -2.57 | n.s. | n.s. |
| Grain (13% moisture) (Mg ha<sup>-1</sup>) | 1.97 | 2.38 | 0.40 | 3.07 | 2.51 | -0.56 | n.s. | n.s. |
| Green leaf area (%) | 95.80% | 85.18% | -10.62% | 98.38% | 75.68% | -22.70% | ** | *** |
| OA (MPa) |  |  | -0.66 |  |  | -1.48 | *** | *** |
| Stomatal density (mm<sup>-2</sup>) | 124.77 | 131.35 | 6.58 | 119.46 | 134.74 | 15.27 | * | * |
| Guard cell length (μM) | 80.17 | 73.95 | 6.22 | 84.03 | 70.15 | 13.89 | * | ** |
| Stomatal index | 20.47 | 21.37 | 0.89 | 18.33 | 16.54 | -1.79 | n.s. | n.s. |
| Leaf 1 length (cm) | 68.65 | 65.43 | -3.41 | 75.91 | 72.30 | -3.61 | * | * |
| C/N | 17.60 | 22.85 | 5.25 | 18.47 | 24.04 | 5.57 | ** | ** |
| N<sub>org</sub> (mg/g DW<sup>-1</sup>) | 24.10 | 18.86 | -4.30 | 23.74 | 18.12 | -4.19 | ** | ** |

<sup>1</sup>Significant *p*-values determined using a one-way ANOVA are denoted by asterisks; \* = *p* < 0.05, \*\* = *p* < 0.005, \*\*\* = *p* < 0.0005, (n.s. = not significant).

Prior to 65 DAP, control and post-flowering drought plots for both genotypes received an equal volume of water once per week, matched to average evapotranspiration rates across the entire field (Fig. 1B-C, see Methods & Materials). After 65 DAP, post-flowering drought (hereafter, “drought”) plots were terminally water deprived (Fig. 1B-C). From the 30 cm to 60 cm depth, plant-available water was 90% depleted by 92 DAP (27 days without water) in droughted plots. From 90 cm to 120 cm depth, water depletion plateaued at ~75% at 105 DAP (40 days without water). Water-deficit stress in droughted plots decreased leaf water and osmotic potentials in both genotypes (Figure 1D-E, Table 1).

Consistent with the strong drought tolerance of these genotypes, grain yields, seed weights, and forage yields were not significantly decreased in droughted plots in either genotype relative to control (Table 1). Green leaf area significantly declined in both genotypes in droughted plots, but the decline was smaller in the BTx642 genotype relative to RTx430, consistent with the stay-green phenotype of BTx642 (Table 1). Also reflecting its stay-green phenotype, BTx642 extracted more soil water in post-flowering drought plots relative to RTx430 (Table S1). Leaf C/N increased in drought in both genotypes driven primarily by decreased organic N (N_org_) content (Table 1). The post-flowering drought response has been described as a competition between maintaining leaf N content and the N demand of developing seeds (Borrell *et al*., 2001). In this context, although N_org_ levels were reduced by drought, N_org_ remained at ~78% of the level of control plants at D4 (40 days without water), highlighting the extent of post-flowering drought tolerance of these genotypes (Table 1).

### Drought depression of mid-afternoon photosynthesis and stomatal opening

Four sampling dates were selected that span the entirety of the water depletion time-course (sampling dates D1-D4, Fig. 1A-C) to model the photosynthetic and metabolic behavior of sorghum during drought in greater detail. The morning measurements [(collected between 9:30 to 11:00) for net photosynthetic rates (*P_n_*), stomatal conductance (*g_s_*), and operating efficiency of PSII in the light (ΦPSII) revealed few statistically significant differences between control and droughted plots (Fig. 2A-C). Two exceptions were a significant difference in *g_s_* between control and drought in BTx642 and between control and drought in ΦPSII in RTx430 at D4 (105 DAP, 40 days without water, Fig. 2B-C). However, the extent to which photosynthetic rates early in the day were unaffected by prolonged water deprivation helps to explain the maintenance of high grain filling rates in post-flowering drought tolerant sorghum.

**Figure 2:**
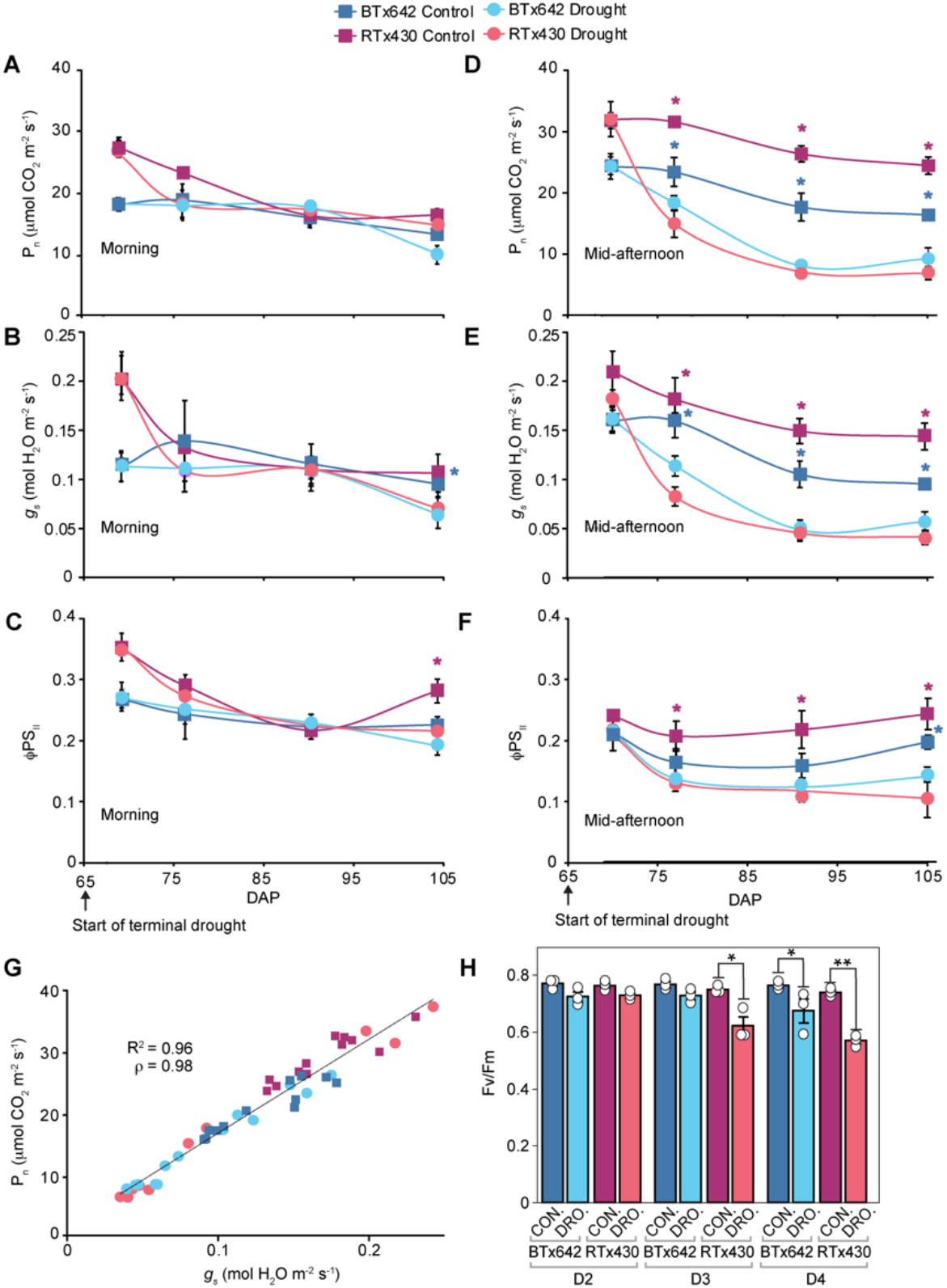
Photosynthetic response to terminal drought stress. (A-H) BTx642 control (dark blue), BTx642 drought (light blue), RTx430 control (purple), and RTx430 drought (pink). (A-C) Measurements in the morning (9:30 to 11:00) and (D-F) collected in mid-afternoon (14:00 to 16:00 on the uppermost three leaves. (A,D) net photosynthetic rate (*P_n_*), (B,E) stomatal conductance (*g_s_*), and (C,F) electron flow through photosystem II (ΦPSII). Light levels ranged between 1651.3 to 1880.5 μmol photons m^−2^ s^−1^ for mid-afternoon measurements and 1001.4 to 1199.4 μmol photons m^−2^ s^−1^ for mid-afternoon measurements. *T_air_* values for D1–D4 were 33.0°C, 39.3°C, 31.6°C, and 31.5°C, respectively, for mid-afternoon measurements and 23.9°C, 28.1°C, 21.6°C, and 25.7°C, respectively, for morning measurements. Mean values ± standard errors (*n* = 3 plots). (A-F) Color of the asterisk denotes which control vs. treatment pair has a *p* < 0.05 by a two-tailed *t*-test. (G) Linear curve fit to mid-afternoon *g_s_* and *P_n_* values for each plot from D1–D4 timepoints with R^2^ and Pearson correlation coefficient (ρ) values. (H) Maximum quantum efficiency of PSII (F_v_/F_m_) measured after 20 min of dark acclimation in the mid-afternoon. Mean values ± standard errors (*n* = 3 plots) with mean values for each individual plot displayed as dots (white). Significant differences, as measured by a two-tailed *t*-test for control vs. treatment pairs, are indicated by asterisks (* < 0.05, ** < 0.005).

In contrast to morning measurements, drought repressed *P_n_, g_s_*, and ΦPSII in both genotypes in mid-afternoon measurements (collected between 14:00 to 16:00) at D2 (77 DAP, 12 days without water), D3 (91 DAP, 26 days without water), and D4 (Fig. 2D-F). *P_n_* and *g_s_* in droughted plots in the morning measurements were either higher or equal to mid-afternoon measurements despite the higher photon flux density in the mid-afternoon at D2, D3, and D4 (Fig. 2A,D, see Table S2 for air temperatures and light levels on sampling dates). This afternoon depression may also act as a drought acclimation strategy to buffer photosynthetic efficiency upon water limitation (Epron *et al*., 1992; Yin *et al*., 2006).

In control plots, mid-afternoon *P_n_* and *g_s_* was higher at all timepoints in RTx430 relative to BTx642 (Fig. 2A,D). Phenotypic comparisons between genotypes grown in separate plots can be made because (a) the total biomass (i.e – forage at 65% moisture, Table 1) was nearly equivalent between genotypes in control plots, (b) both genotypes received the same amount of water (see Materials and Methods), and (c) the comparison between these genotypes grown under equivalent conditions has been made in other published works (Xu *et al*., 2018; Gao *et al*., 2019; Varoquaux *et al*., 2019). That BTx642 maintained more closed stomata in control plots agrees with the lower rates of soil water extraction in BTx642 in irrigated plots relative to RTx430 (Table S2). Thus, well both RTx430 and BTx642 can be considered drought tolerant genotypes given their yield data, their drought tolerance strategies diverge somewhat (Table 1, Fig. 2). BTx642 maintained a constitutively water conservative growth strategy, whereas RTx430 only induced this response in drought. This is also an example of how high photosynthetic rates under well-watered conditions is not predictive of higher photosynthetic rates in drought conditions in the absence of other beneficial traits (Harris *et al*., 2007; Blum, 2009).

*P_n_* and *g_s_* were correlated for all mid-afternoon measurements in both genotypes (Fig. 2G). Along with stomatal closure, photoinhibition can contribute to the depression of *P_n_* in moderate and severe drought. Dark-acclimated maximum quantum efficiency of PSII (F_v_/F_m_) was not significantly depressed in droughted plants until D4 in the stay-green genotype BTx642 and not until D3 in RTx430 (Fig. 2H). Thus, stomatal closure appears to have been the primary cause of drought-induced depression of total daily photosynthetic rates prior to D4 (40 days without water) in the stay-green genotype BTx642 and D3 (26 days without water) in RTx430 (Fig. 2A,D,G-H). Later in the drought period, photoinhibition and loss of green leaf area in the upper canopy also contributed to the depression of total daily photosynthetic rates, particularly in RTx430 (Table 1, Fig. 2H). In fact, the drought-induced decline in F_v_/F_m_ provides a likely explanation for the depression of intrinsic water-use efficiency (WUE_i_) specifically in droughted RTx430 in mid-afternoon measurements (Fig. S2).

A somewhat unexpected observation was that BTx642 and RTx430 droughted plants had small, but significantly increased stomatal density and decreased guard cell length on the abaxial surface of their upper canopy leaves (Table 1). Given that drought was imposed post-anthesis, differences in stomatal density and morphology cannot be explained by the emergence of new leaves with substantial drought-induced developmental changes. However, the absence of a difference in stomatal index (i.e. stomatal density normalized by epidermal cell density) between droughted and control samples suggests that increased stomatal density and reduced guard cell length can be attributed to a reduction in epidermal cell size and the shorter total length of the upper canopy leaves in drought conditions (Clifton-Brown *et al*., 2002).

Drought also depressed the diurnal turnover of transitory photosynthate reserves in leaf tissue (Fig. 3). Starch levels in leaves at dawn were significantly reduced relative to levels at the prior dusk timepoint in both genotypes in control plots at D2, D3, and D4 (Fig. 3A). In contrast, the difference in starch levels between dusk and dawn timepoints was attenuated in droughted plots and fell below the threshold of significance except for BTx642 at D4. A similar trend was present in the leaf sugar content (measured as the sum of leaf sucrose, glucose, and fructose, Fig. 3B). Dawn levels of leaf sugars were significantly lower than dusk at D3 and D4 in both genotypes in control plots and the difference between dusk and dawn levels for total leaf sugars in droughted plots was attenuated and not statistically significant. Dark respiration (*R_d_*) was repressed in both genotypes in drought, providing an explanation for the reduced rate of turnover in transitory photosynthate reserves (Fig. 3C). Yet, the inhibition of diurnal turnover of photosynthate reserves did not impact the final yields, showing that these plants are able to retune their allocation of photosynthate in a manner that does not disrupt grain filling (Table 1). In this light, the inhibition of *R_d_* in droughted plants can represent one means of limiting carbon loss in conditions of restricted photosynthate production (Fig. 2D, 3C) (Ayub *et al*., 2011).

**Figure 3:**
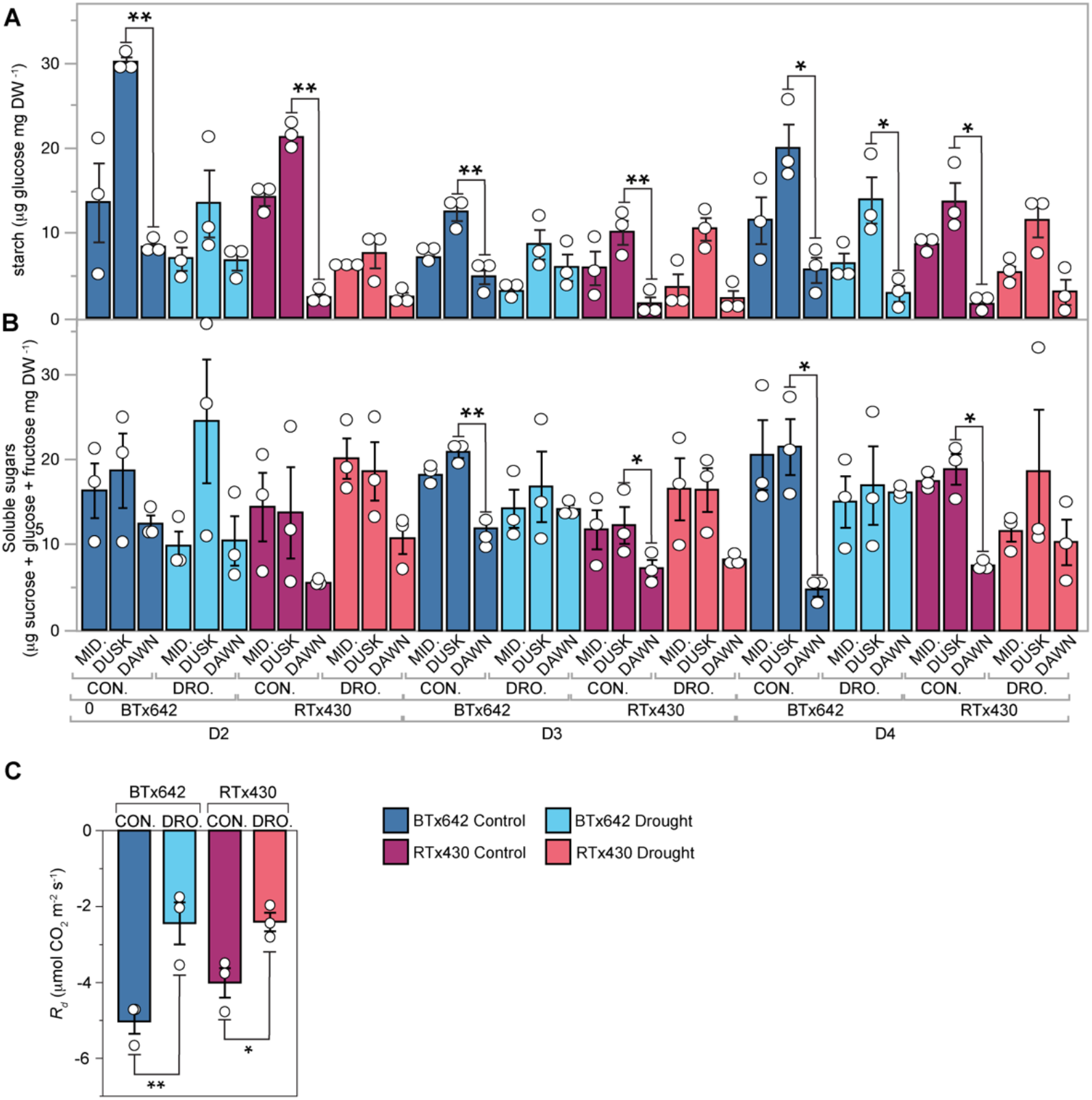
Diurnal starch and sugar accumulation and dark respiration rate. (A) Starch levels quantified as glucose forming units and (B) leaf soluble sugars quantified as the sum of sucrose, glucose, and fructose from the upper canopy (leaves 1-3) collected within an hour of dawn (DAWN), midday (MID.), and within an hour of dusk (DUSK). (C) Dark respiration (*R_d_*) quantified after 20 min of dark acclimation in mid-afternoon. Mean values ± standard errors (*n* = 3 plots) with mean values for each individual plot displayed as dots (white). Significant differences as measured by a two-tailed *t*-test for control vs. treatment pairs are indicated by asterisks (* < 0.05, ** < 0.005). BTx642 control (dark blue), BTx642 drought (light blue), RTx430 control (purple), and RTx430 drought (pink).

### Metabolomic responses to post-flowering drought

Several metabolites linked to pre-flowering drought responses were unresponsive to drought at even the D4 timepoint (40 days of terminal water deprivation). For instance, levels of the drought-responsive hormone abscisic acid (ABA) were not significantly induced in droughted plots except in roots of RTx430 at D2 (Fig. 4A). This suggests that induction of ABA across an entire tissue type, such as roots or leaves, does not factor heavily into the post-flowering drought response in sorghum (Tuberosa *et al*., 1994). Further, ABA is thought to induce biosynthesis of the key osmoprotectant proline in drought and fittingly, proline was not induced by drought in either genotype under these conditions (Fig. 4A) (Ogbaga *et al*., 2014) (Cao *et al*., 2020). These results highlight the uniqueness of the post-flowering drought metabolic response relative to pre-flowering drought responses in sorghum.

**Figure 4:**
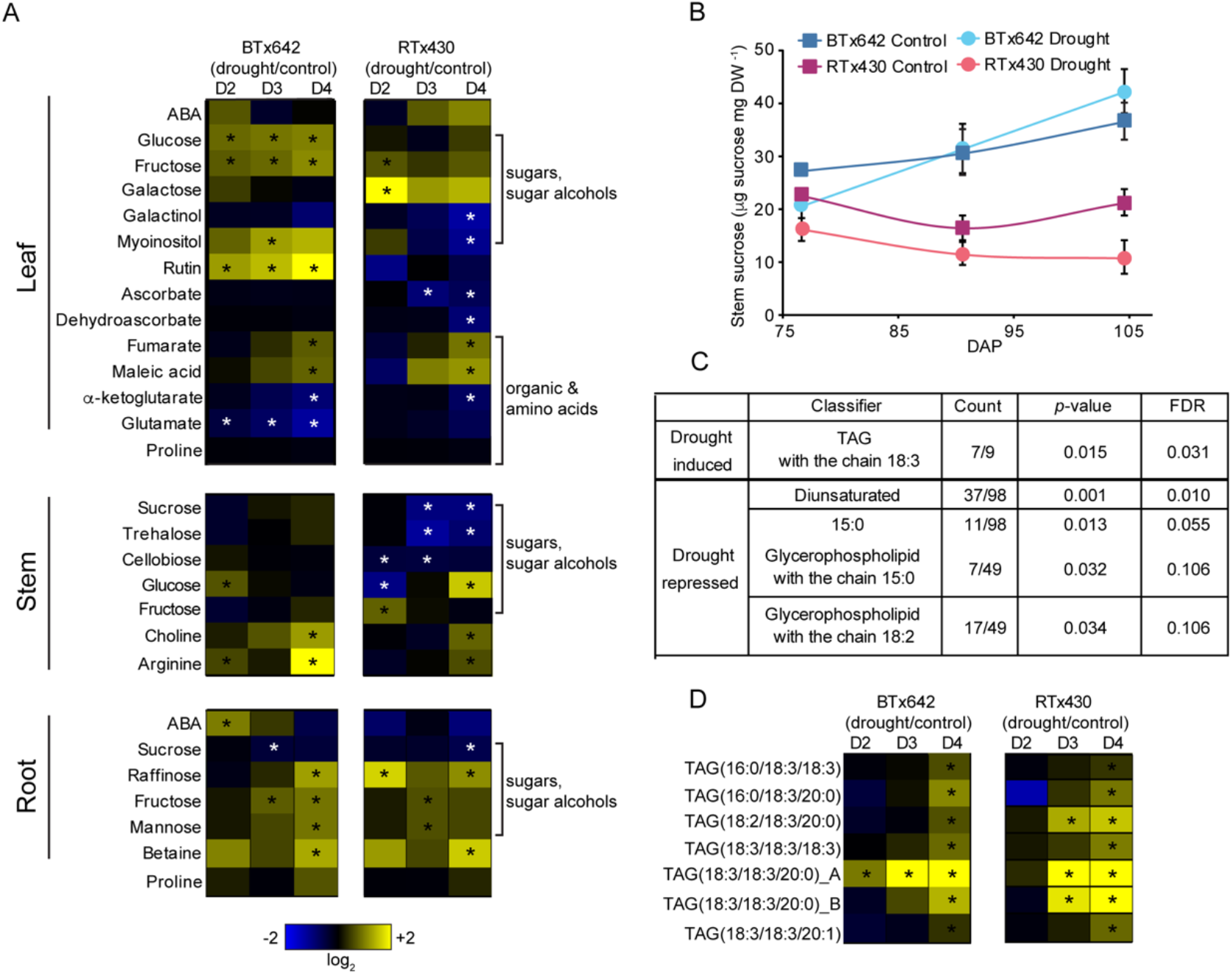
Metabolic and lipidomic acclimation to terminal drought stress in sorghum. (A) Differential abundance for selected metabolites from the three uppermost leaves (upper panels), the upper stem segment below the peduncle (middle panels), and roots in the first 24 cm of soil (lower panels) for BTx642 (left) and RTx430 (right) at three sampling dates: D2 (column 1), D3 (column 2), and D4 (column 3). Metabolite abundance in log_2_ scale with elevated concentration in drought (yellow) and decreased in drought (blue). Significant differences in abundance as determined by a two-tailed *t* test (*p* < 0.05) in drought vs. control are indicated by an asterisk. Black and white asterisks do not have separate meanings. (B) Sucrose levels measured spectrophotometrically from stem samples. Mean values ± standard errors (*n* = 3 plots). BTx642 control (dark blue squares), BTx642 drought (light blue circles), RTx430 control (purple squares), and RTx430 drought (pink circles). (C) Lipid Mini-On (Clair *et al*., 2019) lipid enrichment analysis for drought (BTx642 and RTx430 plots) vs. control (BTx642 and RTx430 plots) leaf samples from the D4 sampling date. Triacylglycerol is abbreviated as TAG in B and C. (D) Differential abundance for leaf TAG lipids with a 18:3 sidechain induced by drought in both genotypes. Visualized in the same manner as panel A.

The two genotypes did induce molecules that can act as osmoprotectants in leaf tissue, albeit different sets of osmoprotectants. Glucose, fructose, and *myo*-inositol were induced in BTx642, whereas RTx430 induced galactose and to a lesser extent fructose as well (Fig. 4A). The induction of leaf osmoprotectants is consistent with the increase in osmotic potential measured at D4 (Figure 1E, Table 1). In contrast to the genotype-specific response of leaf tissue, the osmoprotectant response in root tissue in both genotypes included the shared induction of raffinose, fructose, mannose, and glycine betaine (Fig. 4A).

Levels of the organic acids fumarate and its isomer maleate also increased in droughted leaf tissue in both genotypes, whereas α-ketoglutarate levels decreased (Fig. 4A). Fumarate levels have been negatively correlated with stomatal aperture and are suspected to act as a signal contributing to the regulation of *g_s_* in drought (Araújo *et al*., 2011). The findings here would provide the first indication of a potential role for elevated fumarate levels in signaling drought-induced *g_s_* depression in field-droughted plants (Fig. 2). Decreased α-ketoglutarate levels in both genotypes could be linked to reduced N_org_ levels in droughted plants, as α-ketoglutarate is the organic acid substrate for ammonium assimilation in plants (Fig. 4A, Table 1).

The response of disaccharides to drought in the stem differed strongly between the two genotypes (Fig. 4A,D). A critical component of the functional stay-green trait in sorghum is the prevention of stalk lodging (Rosenow *et al*., 1996; Sanchez *et al*., 2002). Decreased stalk sugar levels during grain filling can contribute to higher stalk lodging rates (Rosenow *et al*., 1996; Sanchez *et al*., 2002; Wang *et al*., 2020). The metabolomic profiling of sugars in the upper stem (below the peduncle) revealed that disaccharide levels (sucrose, trehalose, and cellobiose) all decreased in droughted RTx430 relative to control, whereas they were maintained or slightly increased in the stay-green genotype BTx642 (Fig. 4A,D).

In terms of lipidomics, drought in both genotypes induced levels of polyunsaturated triacylglycerides (TAGs) (Fig. 4B-C). This is particularly interesting because polyunsaturated TAGs have been correlated with the photoprotective response to drought in greenhouse-grown plants of *Arabidopsis thaliana*, wheat, and native Australian grass and tree species (Marchin *et al*., 2017; Ferreira *et al*., 2021). The presence of this response in post-flowering field-droughted plants suggested the activation of photoprotective responses under these conditions, a topic that will be explored further in the next section.

### Stronger photoprotective response minimizes photooxidative stress in BTx642

To maintain photosynthetic leaf area in drought, plants must effectively manage photooxidative stress induced under drought conditions. However, the importance of photoprotective responses to post-flowering drought tolerance has not been examined. In our study, droughted plants of both genotypes did not experience a shared reduction in chlorophyll levels in upper canopy leaves or F_v_/F_m_ until the D4 timepoint, at which point green leaf area had visually declined as well (Fig. 5A, 3H, Table 1). Transcriptomic data revealed that chlorophyllase (*CHL1*), the enzyme responsible for release of phytol during chlorophyll degradation, was induced in both genotypes coinciding with the drop in chlorophyll levels (Fig. S5A). Additionally, q_L_, which represents the fraction of open PSII reaction centers, was not depressed by drought in either genotype until the D3 and D4 timepoints (Fig. S5B). These results highlight the extent to which sorghum minimizes photooxidative stress and damage in post-flowering drought.

**Figure 5:**
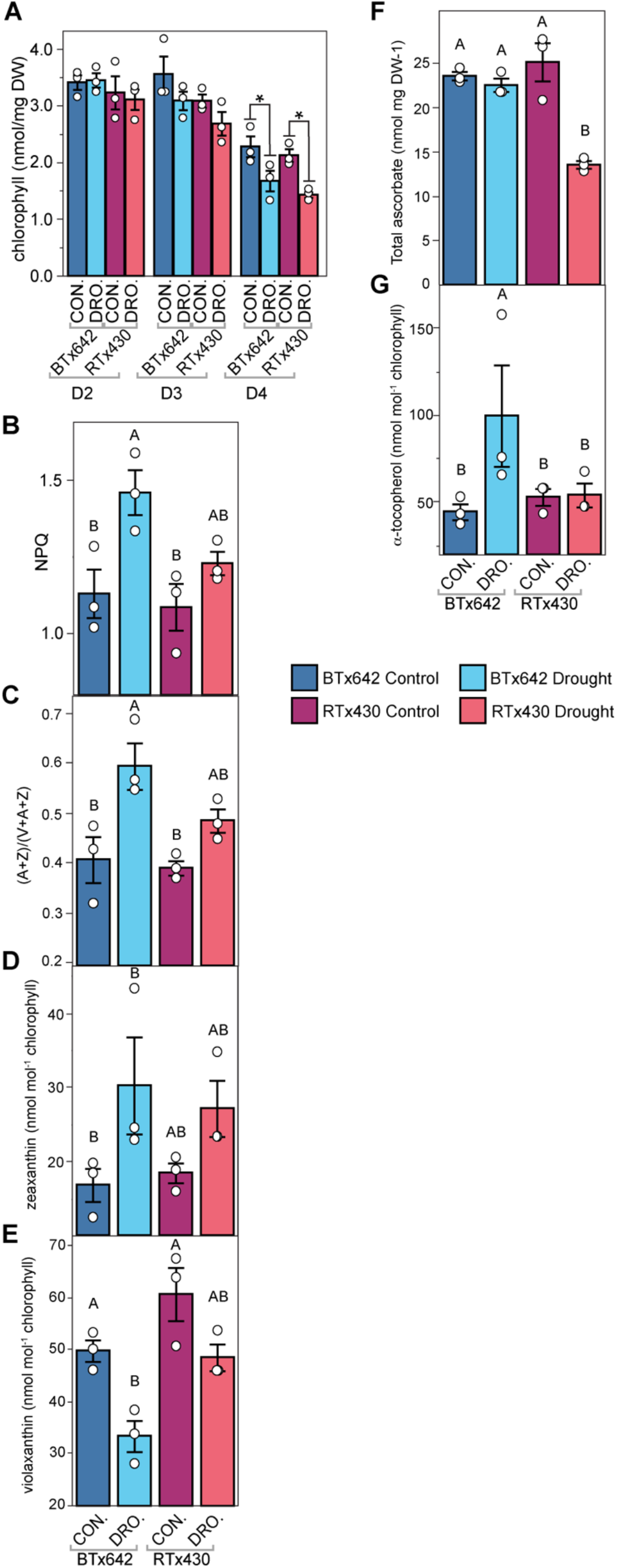
Photoprotective response to terminal drought stress. (A) Chlorophyll levels measured in the uppermost three leaves at the D2–D4 timepoints from samples collected at dawn. From the D4 timepoints, (B) non-photochemical quenching (NPQ) measured at mid-afternoon, (C) epoxidation state of the Violaxanthin + Antheraxanthin + Zeaxanthin (VAZ) pool measured as (A+Z)/(V+A+Z), (D) zeaxanthin, (E) antheraxanthin, (F) total ascorbate levels, and (G) α-tocopherol levels. (C-G) From leaf samples collected in the mid-afternoon. Mean values ± standard errors (*n* = 3 plots) with mean values for each individual plot displayed as dots (white). BTx642 control (dark blue), BTx642 drought (light blue), RTx430 control (purple), and RTx430 drought (pink). (A) Significant differences as measured by a two-tailed *t*-test for control vs. treatment pairs are indicated by asterisks (* < 0.05). (B-G) Mean values that share the same letters are not statistically different, and those that do not share the same letters are statistically different based on one-way ANOVA and post hoc Tukey–Kramer HSD tests.

However, indicators of photooxidative stress tended to emerge sooner and more strongly in droughted RTx430 plants relative to the stay-green genotype BTx642. F_v_/F_m_ in droughted RTx430 was lower at D3 and D4 relative to droughted BTx642 (Fig. 3H). A pattern of greater photooxidative stress in droughted RTx430 relative to BTx642 is also consistent with the observed greater decrease in green leaf area in droughted RTx430 relative to BTx642 (Table 1).

Several lines of evidence point to a more robust drought-induced photoprotective response in the stay-green BTx642 genotype. Non-photochemical quenching (NPQ) was induced specifically in droughted BTx642 in mid-afternoon measurements (Fig. 5B). Supporting this genotype-specific induction of NPQ, the de-epoxidation state of the xanthophyll pool was specifically higher in droughted BTx642 relative to the control conditions (5C-E). These NPQ measurements were performed specifically on non-senesced portions of leaves to maintain an accurate representation of photoprotective responses in tissues that are still active in photosynthesis. Beyond NPQ, higher total ascorbate levels were maintained in droughted BTx642 in contrast to RTx430 (Fig. 4A, 5F). In the transcriptomic data, several ascorbate enzymes were induced in both genotypes in drought (Fig. S4), consistent with the enhanced demand for ascorbate to ameliorate ROS stress (Laxa *et al*., 2019). Further, the chloroplast-localized antioxidant α-tocopherol and the epidermis-enriched photoprotective flavonoid, rutin, were specifically induced in droughted BTx642 (Fig. 5G, 2A). An explanation for why tocopherols might have been induced specifically in droughted BTx642 can be found in the higher transcript level for several genes involved in tocopherol biosynthesis, such as Sobic.004G024600 (*LIL3*), Sobic.010G207900 (*VTE2-2*), and Sobic.006G260800 (*VTE5*), in droughted BTx642 relative to droughted RTx430 (Fig. S6).

Taken together, a stronger photoprotective capacity in droughted BTx642 may limit photooxidative damage and thereby, minimize the extent of drought-induced early leaf senescence under these conditions relative to RTx430 (Table 1). The role of photoprotection in preventing drought-induced leaf senescence is well-established, particularly in perennials (Murchie *et al*., 1999; Munné-Bosch *et al*., 2001; Munné-Bosch and Peñuelas, 2003; Demmig-Adams and Adams, 2006; Challabathula *et al*., 2018). Thus, if future research confirms that stronger photoprotection contributes to the stay-green phenotype in sorghum, then this trait would follow the framework of stay-green as an example of perennial-like traits emerging in an annual plant (Thomas and Howarth, 2000).

### Identifying putative candidate genes underlying stay-green in sorghum

We have combined in-field physiological analysis with transcriptomic and metabolomic analysis of rapidly frozen tissues to reveal new insights into drought tolerance in sorghum. The RTx430 and BTx642 sorghum genotypes maintained *P_n_* values on par with control plants early in the day throughout the grain filling period despite long-term water deprivation (Fig. 1,2). Mid-afternoon depressions in *P_n_* in droughted plants were driven largely by stomatal closure early in the drought period (Fig. 2). In the later stages of drought, levels of photoinhibition increased (Fig. 2H) and green leaf area declined (Table 1) particularly in the RTx430 genotype. The stay-green BTx642 genotype more strongly induced photoprotective responses (Table 1, Fig. 2, 4, 5). These included genotype-specific drought induction of NPQ and tocopherols in BTx642, supporting a previously uncharacterized role for photoprotective pathways in the functional stay-green trait (Fig. 5). Of particular interest from the metabolomic datasets are molecules that may act as regulators of drought tolerance, such as fumarate, which may contribute to drought-induction of stomatal closure, and polyunsaturated TAGs, which may act to minimize excess ROS accumulation (Fig. 4) (Araújo *et al*., 2011; Ferreira *et al*., 2021). In contrast, whole-tissue ABA levels and proline remained largely non-drought responsive (Fig. 4).

One means to alleviate the drought-induced decline in photosynthetic rates observed here in the mid-afternoon would be to attenuate expression or altogether disrupt certain genes that participate in drought-induced stomatal closure in sorghum via gene editing. Such an approach would rest on the idea that genotypes may be overly responsive to the threat of excess water loss in drought. By tuning down this response—but not entirely blocking it—it may be possible to sustain higher photosynthetic rates in the grain-filling period without overly exacerbating water loss rates. Given the stronger response of *g_s_* in RTx430 to drought (Fig. 2), one means to identify sorghum genes that control drought-induced stomatal closure is to look for orthologs to known positive regulators of stomatal closure, as characterized in model plant species, and to determine which are more strongly induced by drought in RTx430 relative to BTx642 (Khan *et al*., 2013; Ge *et al*., 2015). Examples that fit this pattern include the MAP kinase Sobic.007G046100 (*MPK4*) and the homologs of ABA-insensitive G-protein α-subunits Sobic.003G242200, Sobic.009G213000, and Sobic.003G198200 (Fig. 7). As sorghum cell-type specific transcriptomes are made available, it will be important to determine which of these putative stomatal regulators are strongly expressed in guard cells.

Drought also causes photooxidative stress, leading to ROS-induced early leaf senescence. In this study, BTx642 appears to have stronger photoprotective capacity in post-flowering drought based on higher NPQ levels (Fig. 5B-C), induction of photoprotective molecules (e.g. α-tocopherol (Fig. 5D) and rutin (Fig. 4)), maintenance of high ascorbate pool size (Fig. 6), and less photooxidative damage and leaf senescence relative to RTx430 (Table 1, Fig. 2). Given the higher abundance of transcripts for tocopherol biosynthetic genes in the stay-green genotype BTx642, boosting tocopherol levels in RTx430 via over-expression of tocopherol biosynthesis enzymes may be one avenue to inhibit drought-induced early leaf senescence in sorghum (Liu *et al*., 2008; Zhan *et al*., 2019). A second avenue to improve stay-green capacity in RTx430 could involve increasing NPQ capacity by post-flowering drought over-expression of the NPQ regulator *PSBS*, or by supporting a more de-epoxidized xanthophyll cycle pool for NPQ (Głowacka *et al*., 2018).

**Figure 6:**
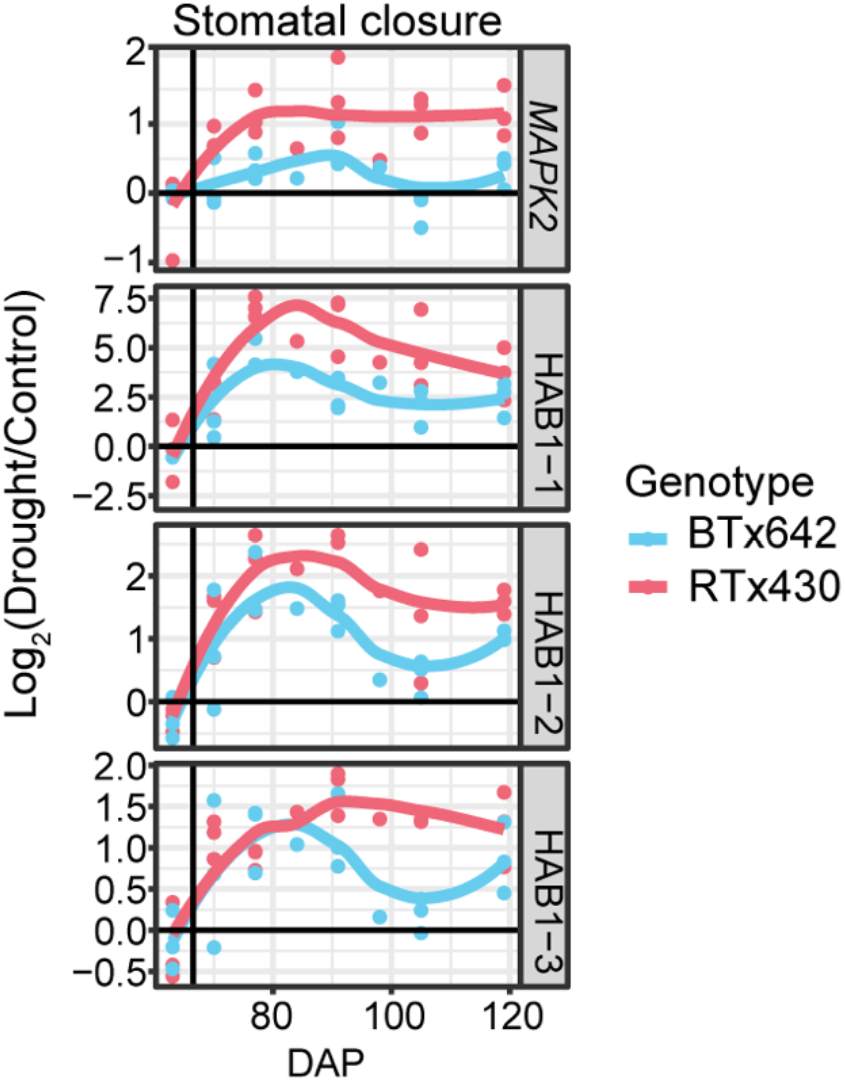
Log_2_ fold change in transcript abundance in leaf tissue for drought versus control for the orthologs of drought-responsive regulators of stomatal closure that are induced more strongly in RTx430 in response to post-flowering drought, Sobic.007G046100 (*MAPK2*), Sobic.003G242200 (*HAB1-1*), Sobic.009G213000 (*HAB1-2*), and Sobic.003G198200 (*HAB1-3*). BTx642 in light blue, RTx430 in pink.

As the challenges of climate change impact agriculture in the coming decades, the ability to confer a functional post-flowering drought tolerance to drought-susceptible genotypes provides a path towards maintaining or even improving yields with limited water inputs. The diverse, yet synergistic, pathways underlying sorghum drought tolerance that are outlined in this work support several possible avenues to achieve this goal, with field-level resolution substantiating these phenotypes. Building upon this study, our insights into the transcriptional and molecular underpinnings of stay-green in sorghum can be used to select targets for gene editing to test their involvement in post-flowering drought tolerance and to improve crop productivity under drought.

## Abbreviations

WUEi: Intrinsic water-use efficiency
ROS: Reactive oxygen species
NPQ: Non-photochemical quenching
DAP: Days after planting
F_v_/F_m_: Dark-acclimated maximum quantum efficiency of PSII
*P_n_*: Net assimilation of CO_2_ in the light
*g_s_*: Stomatal conductance
ΦPSII: Operating efficiency of PSII in the light
SPE-IMS-MS: Ion mobility spectrometry and mass spectrometry
ABA: Abscisic acid
TAG: Triacylglycerides
*R_d_*: Respiration in the dark

## Acknowledgments

We thank for their help in collecting field data and processing plant samples, Mimi Broderson, Ryan McCombs, Olga Gaidarenko, Christine Pagotan, Gauri Kapse, Chandler Sutherland, Victoria Kim, Kiflom Aregawi, Claudia Jane Bucheli, Rachel Bosynak, Lili Montaya, Cynthia Amstutz, Gabriella Benko, Jeffrey Johnson, Jianqiang Shen, and Chase Turnbull. We thank Nicholas Karavolias for his advice on data analysis.

## Author Contributions

CRB, KKN, PGL, JD, RBH, and KK designed and planned the research. CRB and DP conducted experiments, collected field samples, and analyzed data. LGC, OD, and AK conducted experiments and collected/processed field samples. RBH, BJC, AB, JYL, YMK, JEK, KJB, and VP conducted experiments and analyzed data. CA, JP, and JS collected/processed field samples. CRB wrote the manuscript with contributions from BJC, AB, JYL, and YMK. The manuscript was edited by KKN, JD, PGL, DP, and AK. OD and AK contributed equally to this work. CA, JP, and JS contributed equally to this work. AB, JYL, and YMK contributed equally to this work.

## Data availability

The data that support the findings of this study are available from the corresponding author upon reasonable request.

## Tables

**Table S1:**
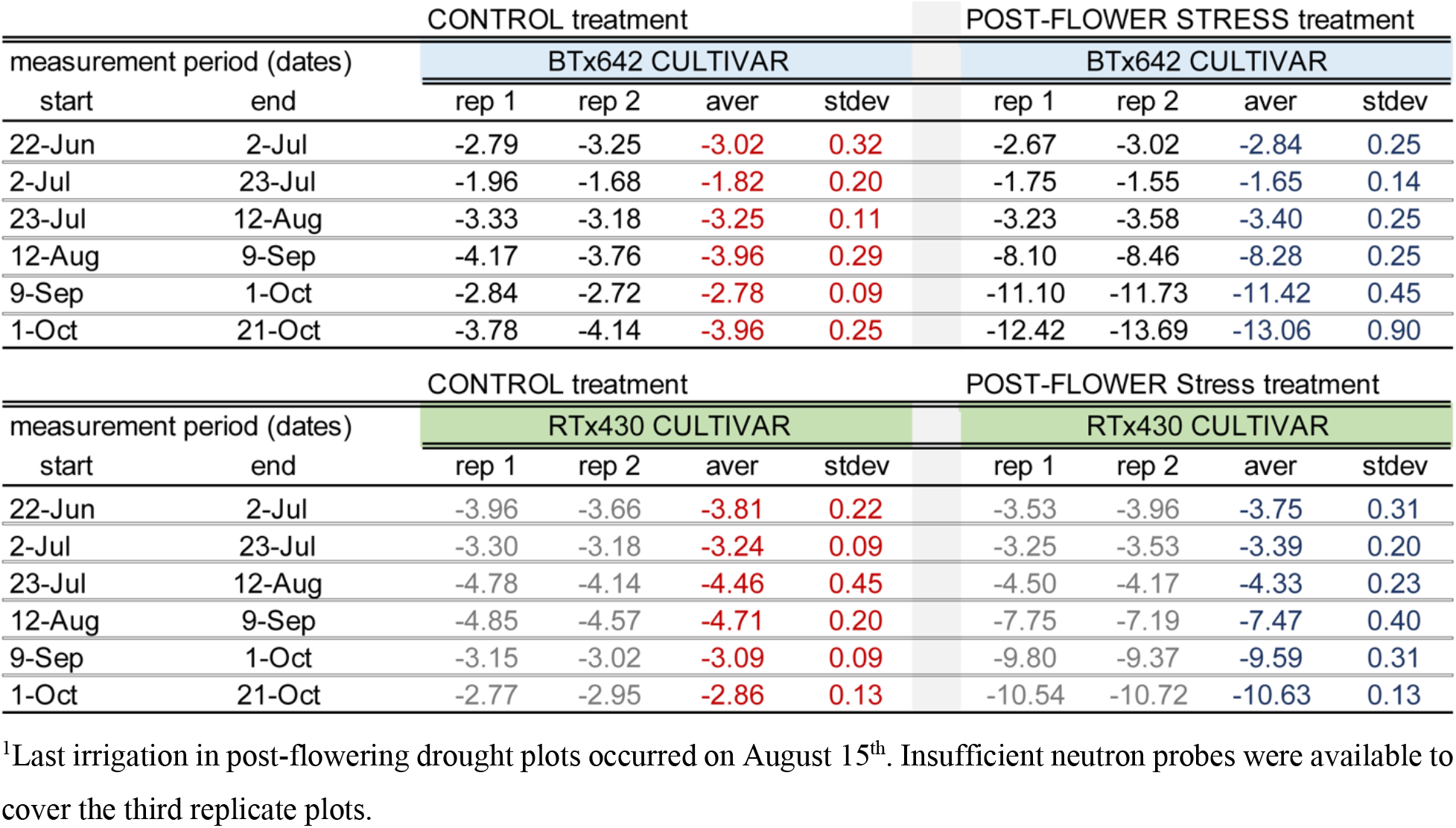
Change in total soil profile water content(cm soil water) during time periods shown

**Table S2:** Ambient light and air temperature at four sampling trips

|  | Morning |  | Mid-afternoon |  |
| --- | --- | --- | --- | --- |
| Sampling trip | Avg. temp(°C) | Light intensity ($\mu\text{mol m}^{-2} \text{s}^{-1}$) | Avg. temp(°C) | Light intensity ($\mu\text{mol m}^{-2} \text{s}^{-1}$) |
| D1 | 24.26 | 998.03 | 32.90 | 1444.15 |
| D2 | 28.30 | 1199.99 | 39.14 | 1879.52 |
| D3 | 21.58 | 1050.10 | 31.73 | 1649.98 |
| D4 | 26.00 | 1050.60 | 31.49 | 1591.29 |

## Figures

**Supplemental figure 1:**
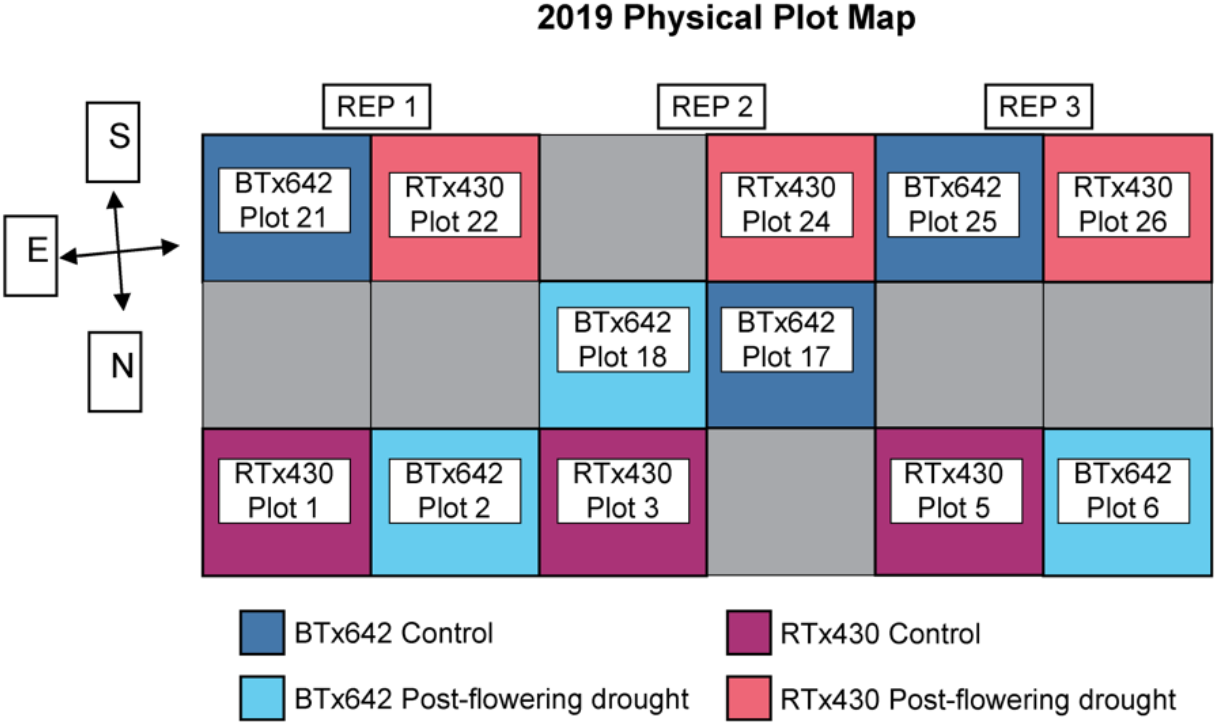
Field layout of 2019 sorghum field trials. Sorghum grown in sandy loam soils at the University of California Kearney Agricultural Research and Extension Center in Parlier, CA, USA. BTx642 control (dark blue), BTx642 drought (light blue), RTx430 control (purple), and RTx430 drought (pink).

**Supplemental figure 2:**
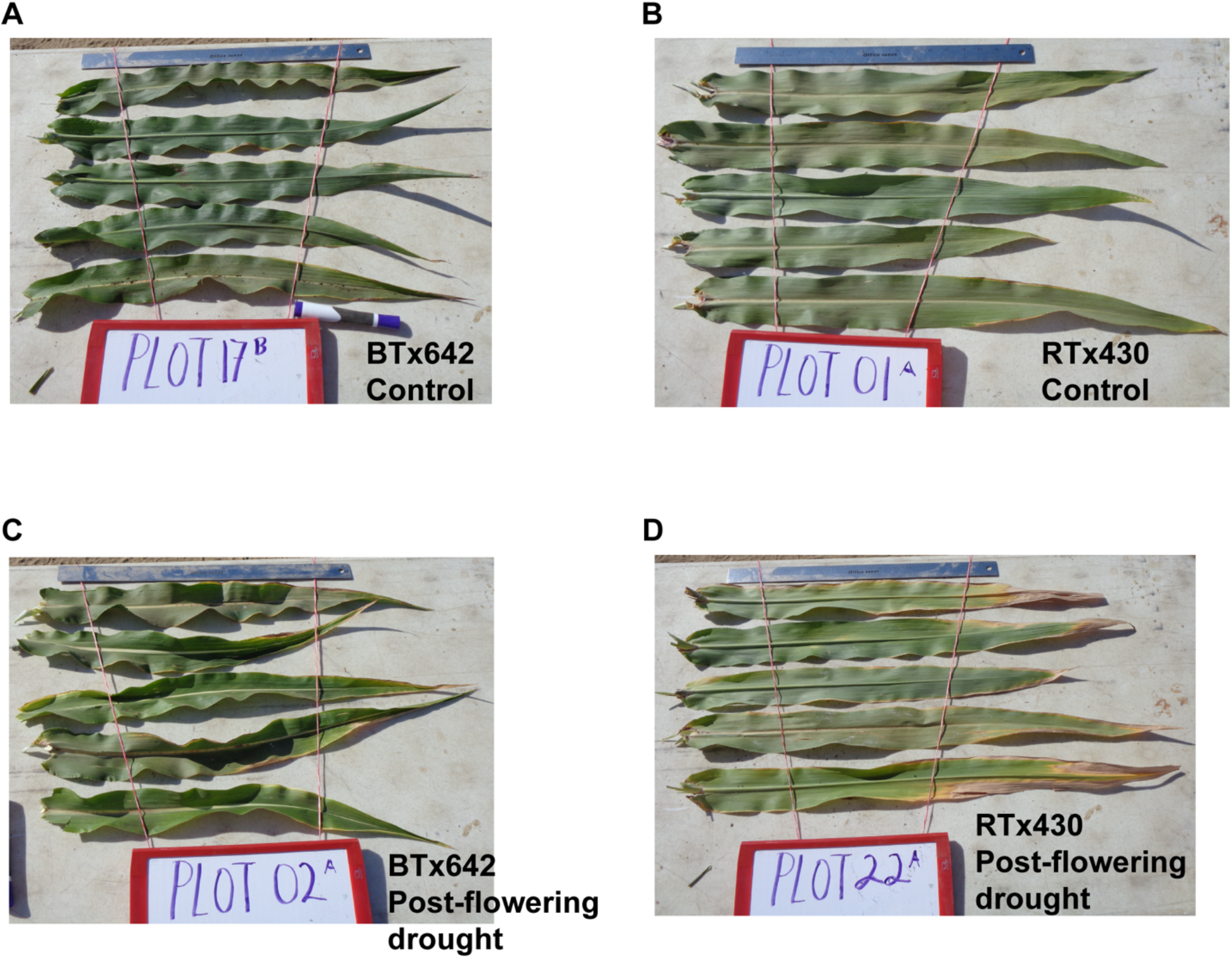
Representative photos of leaves in control and post-flowering drought plots. Leaves sampled on D4 (40 days without water for droughted plots) from five different plants and a randomly selected leaf from the uppermost three leaves including flag leaves. (A) BTx642 control, (B) RTx430 control, (C) BTx642 post-flowering drought, (D) RTx430 post-flowering drought.

**Supplemental Figure S3:**
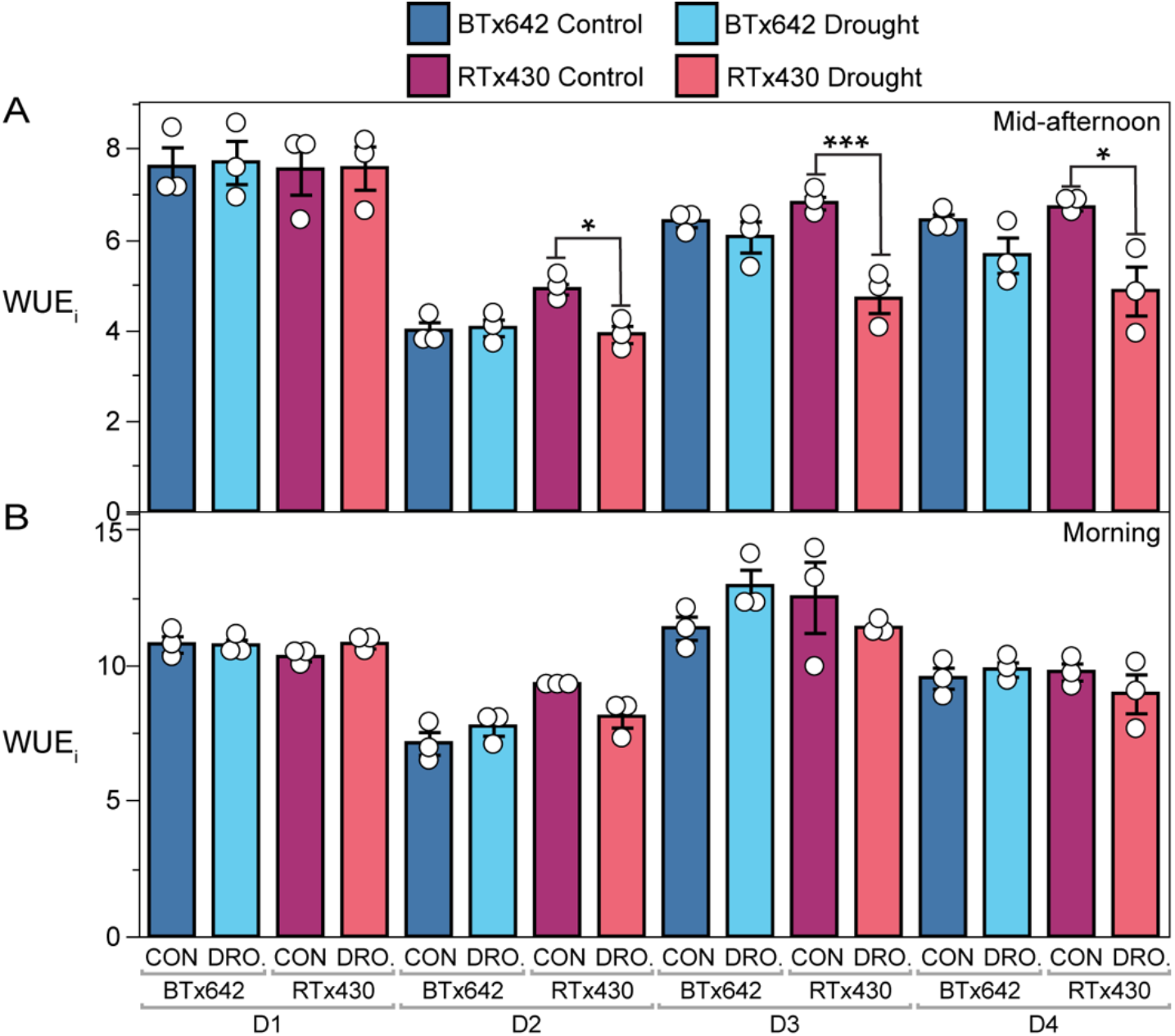
Instantaneous water-use efficiency (WUE_i_) for (A) mid-afternoon measurements (14:00 to 16:00) and (B) morning measurements (9:30 to 11:00). Mean values ± standard errors (*n* = 3 plots) with mean values for each individual plot displayed as dots (white). Significant differences as measured by a two-tailed *t*-test for control vs. treatment pairs are indicated by asterisks (* < 0.05, ** < 0.005, *** < 0.0005). BTx642 control (dark blue), BTx642 drought (light blue), RTx430 control (purple), and RTx430 drought (pink).

**Supplemental figure 4:**
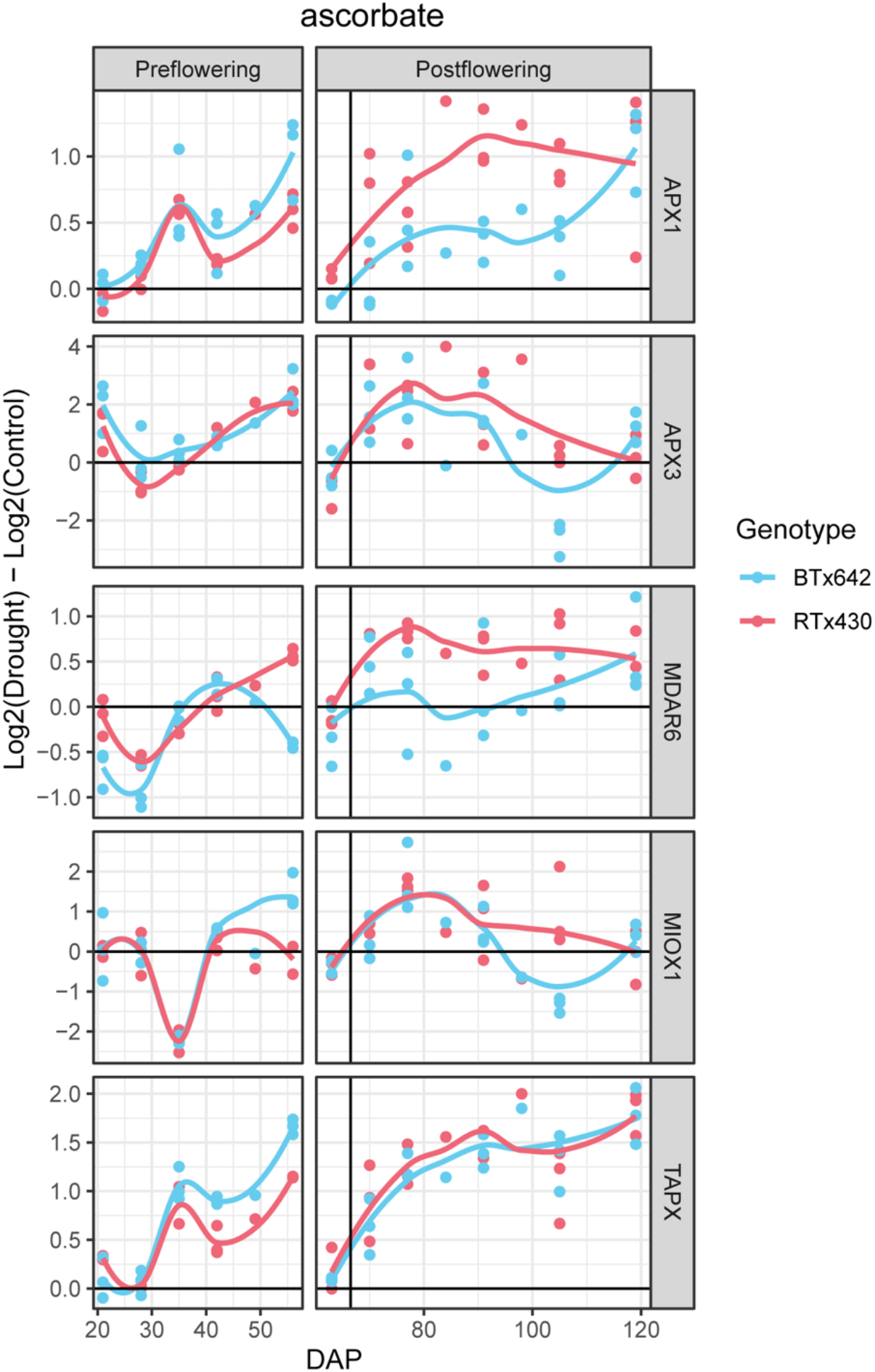
Log_2_ fold change in transcript abundance in drought versus control in leaf tissue for ascorbate peroxidase genes, Sobic.001G410200 (*APX1*), Sobic.006G021100 (*APX3*), Sobic.006G084400 (*TAPX*); for monodehydroascorbate reductase gene, Sobic.007G038600 (*MDAR6*). BTx642 in light blue, RTx430 in pink.

**Supplemental figure 5:**
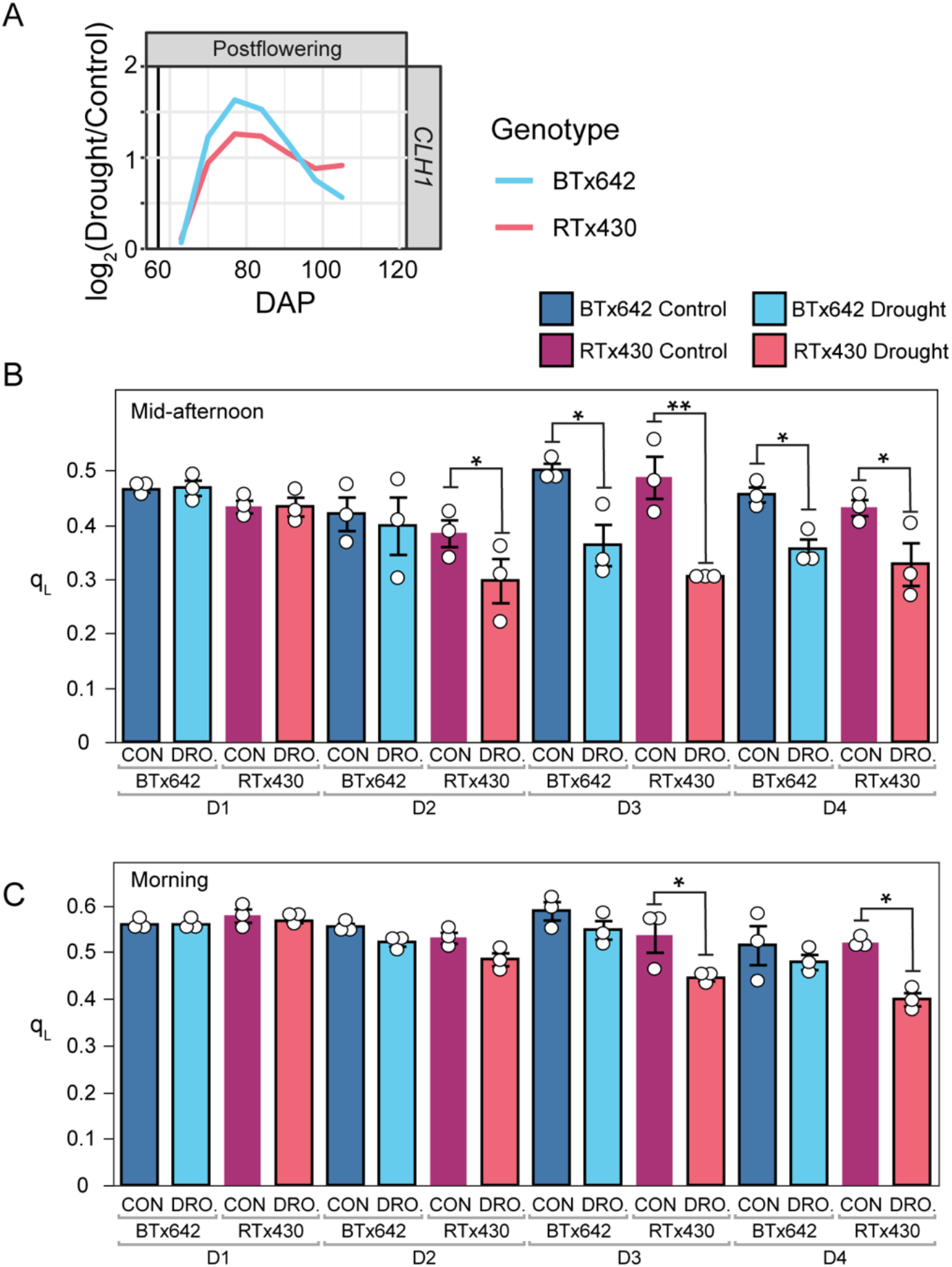
Induction of chlorophyll degradation and response of q_L_ in terminal drought. (A) Log_2_ fold change in transcript abundance in leaf tissue for drought versus control for the chlorophyllase gene, Sobic.007G168000 (*CHL1*). (B-C) Redox state of the fraction of open PSII reaction centers measured as q_L_ for (B) mid-afternoon measurements (14:00 to 16:00) and (C) morning measurements (9:30 to 11:00). Mean values ± standard errors (*n* = 3 plots) with mean values for each individual plot displayed as dots (white). Significant differences as measured by a two-tailed *t*-test for control vs. treatment pairs are indicated by asterisks (* < 0.05, ** < 0.005). BTx642 control (dark blue), BTx642 drought (light blue), RTx430 control (purple), and RTx430 drought (pink).

**Supplemental figure 6:**
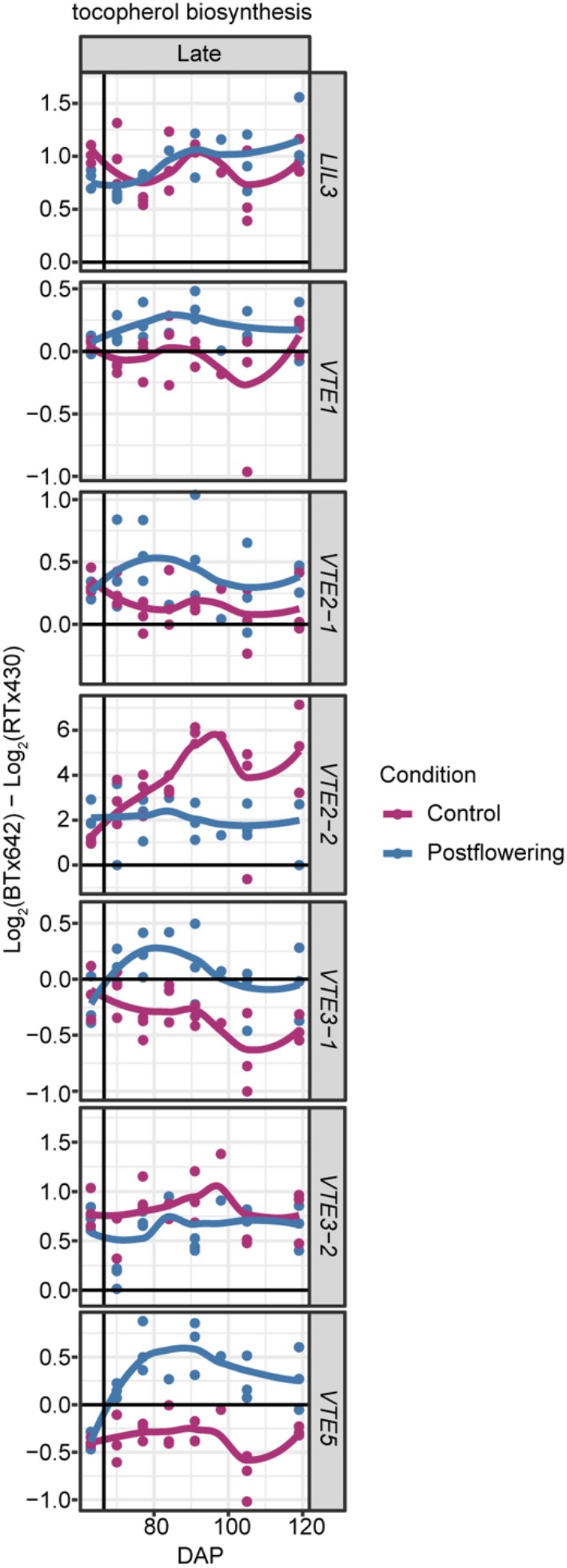
Log_2_ fold change in transcript abundance across genotypes (BTx642 versus RTx430) in leaf tissue for drought versus control for tocopherol biosynthesis genes: Sobic.004G024600 (*LIL3*), Sobic.004G125800 (*VTE1*), Sobic.010G215600 (*VTE2-1*), Sobic.010G207900 (*VTE2-2*), Sobic.008G171300 (*VTE3-1*), Sobic.008G171000 (*VTE3-2*), and Sobic.006G260800 (*VTE5*). Log_2_ (BTx642 control/RTx430 control) in dark blue and log_2_ (BTx642 post-flowering drought/RTx430 post-flowering drought) in purple.

## Supplemental materials

### Supplemental materials & methods

#### Field growth conditions and crop evapotranspiration

Crop evapotranspiration was determined using potential evapotranspiration measured at an on-site CIMIS (CA Irrigation Management Information System) weather station multiplied by the crop coefficient, which was adjusted according to crop growth stage. See Xu et al. for details regarding estimation of evapotranspiration rates, irrigation management, and drought treatment measurements (Xu *et al*., 2018). Briefly, Crop Water Stress Index measurements were performed using a combination of fixed position infrared thermometers (IRT) and handheld-IRT mid-afternoon measurements on select dates across drought progression in combination with continuous monitoring of air temperature and relative humidity using an in-field weather station (O’Shaughnessy *et al*., 2012). Soil water potential and soil water content were monitored using soil matric potential sensors and neutron probes, respectively. Matric potential sensors will be installed at multiple depths to represent the most active water uptake portions of the crop root zone (at 30 cm increments from 15 cm depth to 105 cm). Neutron probe access tubes were installed to a depth of 1.5 m in select plots, and soil water measurements were taken at 30 cm increments.

The treatment conditions were assigned via a randomized block design, where the fields were divided into 18 plots of 10 rows each, each plot randomly assigned a watering treatment (control and post-flowering drought) and in 2019, three genotypes (RTx430, BTx642, and RTx7000), with 3 replicates, for a total of 18 plots. RTx7000 plots were sampled only sparingly in 2019 due to low seed germination rates as a consequence of non-uniform seed quality (grey boxes in Fig. S1).

#### Leaf phenotypic traits

To avoid edge effects, all phenotypic measurements were conducted on plants in the interior of each plot. Leaf water potential (ψ_I_) was measured and flash-frozen samples collected for measuring osmotic potential (ψ_s_) readings in the mid-afternoon at the end of sampling day D4 (on September 24, 2019) using PSY1 leaf psychrometers (ICT International, Armidale, Australia) and carefully following the instructional protocols provided with these instruments. Control plots had not received water for six days and post-flowering drought plots had not received water for 41 days. Three plants per plot were selected and the uppermost leaf below the flag leaf of the main culm was measured and samples collected at the leaf midpoint (avoiding midrib, equal distance from the leaf tip and base). The waxy surface of the leaf was partly removed before measuring ψ_I_ by gently rubbing the measurement area with aluminum oxide (642991, Sigma-Aldrich, St. Louis, MO, USA) and then rinsed to remove the excess aluminum oxide before measurement. Stable ψ_I_ and ψ_s_ values were reached within 40 min of initiating the measurement. Green leaf area and relative water content (RWC) samples were also collected on September 24, 2019. Green leaf area was determined by imaging the three uppermost leaves of ten randomly selected plants per plot and area quantification of visibly senesced and non-senesced leaf area using ImageJ (Schindelin et al., 2012). A ruler was used in each photograph to normalize the leaf size between digital photographs.

Steady-state photosynthetic rates (P_n_), stomatal conductance (*g_s_*), photosystem II operating efficiency (ΦPSII), fraction of closed PSII reaction centers (q_L_), PSII maximum photochemical efficiency in the light (F_v_’/F_m_’), instantaneous water use efficiency (WUE_i_), and intracellular CO_2_ / atmospheric CO_2_ (C_i_/C_a_) were determined using LI-COR 6400XT infrared gas analyzers with a chlorophyll fluorometer attachment (LI-COR, Lincoln, NE, USA). Light levels (10% blue light) and chamber temperature was matched to ambient conditions. q_L_ was calculated as (1/F_s_ - 1/F_m_’)/(1/F_o_’ - 1/F_m_’), ΦPSII as (F_m_’ - F_s_)/F_m_’, and F_v_’/F_m_’ as (F_m_’ - F_o_’)/F_m_’. Respiration in the dark (R_d_) and the maximum photochemical efficiency of PSII in the dark (F_v_/F_m_) were measured following a 20-min dark acclimation of attached leaves in the field in the mid-afternoon on sampling date D4 (09/23 - 09/24/2019). Block temperature was again matched to ambient temperatures and relative humidity held between 50% to 60%. Non-photochemical quenching (NPQ) was measured as (F_m_ – F_m_’)/F_m_’ on the same dark-acclimated leaves following a 10-min actinic light exposure (10% blue / 90% red) where the light level was matched to the ambient light level measured prior to this set of measurements (1650 μmol photons m^−2^ s^−1^).

Stomatal density and guard cell length were quantified using ImageJ from light microscopy images of leaf peels collected on the D4 sampling day from the abaxial leaf surface as described in Lopez et al. (Lopez *et al*., 2017).

#### Metabolite extraction and quantification

Harvested and frozen root, stem, and leaf tissue were ground in a cryogenic Freezer Mill (SPEX SamplePrep 6875D, Metuchen NJ USA) for 2-3 cycles of 2-3 min, with 1 min cooling in between. Samples were then stored at −80°C. Chlorophyll and total ascorbate were extracted and quantified via spectrophotometry as previously described (Arnon, 1949; Queval and Noctor, 2007). Carotenoids and tocopherol were also extracted using acetone and quantified by high-performance liquid chromatography using standard protocols as previously described (Müller-Moulé *et al*., 2002). Soluble sugars were extracted using ethanol and starch in the pellet was solubilized by an amylase/amyloglucosidase treatment and quantified spectrophotometrically using standard protocols as previously described (Stitt *et al*., 1989; Smith and Zeeman, 2006). Metabolites for gas chromatography-mass spectrometry/mass spectrometry (GC-MS), lipidomics, and ion-mobility spectroscopy (IMS) were extracted using a methanol-chloroform extraction as previously described (Handakumbura *et al*., 2017). Leaf tissue samples from sampling days D2, D3, and D4 were analyzed by GC-MS, lipidomics, and IMS. Metabolomic data was collected for stem and root samples from D2, D3, and D4 sampling dates exclusively by IMS.

#### Metabolomics using GC-MS

The flash-frozen leaf and root were mechanically ground separately using a cryogenic freezer mill (SPEX, Metuchen, NJ) kept at cryogenic temperatures with liquid nitrogen. Then MPLEx extraction was applied to the samples which were weighed at 1 g (Nakayasu *et al*., 2016). Then, the samples were completely dried under a speed vacuum concentrator. The dried metabolites were chemically derivatized and analyzed by gas chromatography-mass spectrometry or GC-MS as reported previously (Kim *et al*., 2015). Briefly, dried samples were derivatized by adding 20 μL of methoxyamine solution (30 mg/mL in pyridine) and were incubated at 37 °C for 90 min to protect the carbonyl groups and reduce carbohydrate isoforms. Then, 80 μL of N-methyl-N-(trimethylsilyl)-trifluoroacetamide with 1% trimethylchlorosilane was added to each sample and incubated for 30 min as a minimum. The derivatized samples were analyzed by GC/MS within 24 hours after the derivatization. Data collected by GC/MS were processed using the Metabolite Detector software, version 2.5 beta (Hiller *et al*., 2009). Retention indices of detected metabolites were calculated based on analysis of the fatty acid methyl esters mixture (C8 - C28), followed by chromatographic alignment across all analyses after deconvolution. The intensity values of selected three fragmented ions after deconvolution were integrated for a peak value of metabolite. Metabolites were initially identified by matching experimental spectra to a PNNL augmented version of the Agilent Fiehn Metabolomics Library containing spectra and validated retention indices for almost 900 metabolites (Kind *et al*., 2009) and additionally cross-checked by matching with NIST14 GC/MS Spectral Library. All metabolite identifications were manually validated to minimize deconvolution and identification errors during the automated data processing. The data were log_2_ transformed and then mean-centered across the log_2_ distribution.

#### Lipidomics

Total lipid extracts (TLEs) were analyzed as outlined in Kyle et al. (2017). Briefly, a Waters Acquity UPLC H class system interfaced with a Velos-ETD Orbitrap mass spectrometer was used for LC-ESI-MS/MS analyses. 10 μL of the reconstituted sample was injected onto a Waters CSH column (3.0 mm x 150 mm x 1.7 μm particle size) and separated over a 34-min gradient (mobile phase A: ACN/H2O (40:60) containing 10 mM ammonium acetate; mobile phase B: ACN/IPA (10:90) containing 10 mM ammonium acetate) at a flow rate of 250 μL/min. TLEs were analyzed in both positive and negative electrospray ionization modes, and lipids were fragmented using alternating higher-energy collision dissociation (HCD) and collision-induced dissociation (CID) (Kyle *et al*., 2017). Identifications were made using LIQUID (Kyle *et al*., 2017) and manually validated by examining the MS/MS spectra for fragment ions characteristic of the classes and acyl chain compositions of the identified lipids. In addition, the precursor ion isotopic profile extracted ion chromatogram, and mass measurement error along with the elution time was evaluated. All LC-MS/MS data were aligned and gap-filled to this target library for feature identification using MZmine 2 (Pluskal *et al*., 2010) based on the identified lipid name, observed m/z, and retention time. Data from each ionization mode were aligned and gap-filled separately. Aligned features were manually verified and peak apex intensity values were exported for statistical analysis.

#### SPE-IMS-MS Metabolomics Analysis

Plant extracts were analyzed by SPE-IMS-MS using a RapidFire 365 (Zhang *et al*., 2016) coupled with an Agilent 6560 Ion Mobility QTOF MS system (Agilent Technologies, Santa Clara, CA, USA). The samples were loaded onto three different SPE cartridges using a 10 μL loop. For the Graphitic Carbon cartridge, the loading solvent consisted of 0.1% formic acid and 99.9% water. The analytes were eluted off the cartridge using a combination of 0.1% formic acid, 49.95% water, 24.98% acetonitrile, and 24.98% acetone. The C18 cartridge used the same loading solvent as the Graphitic Carbon but was eluted using 49.95% methanol, 49.95% IPA, and 0.1% formic acid. The HILIC cartridge was loaded using 90% acetonitrile and 10% 20 mM ammonium acetate and eluted with 90% 20 mM ammonium acetate and 10% acetonitrile. Samples were loaded onto the cartridges at a flow rate of 1.5 mL/min and eluted at a flow rate of 0.6 mL/min. The sample injection parameters were as follows: aspiration time, load time, and elution time were 0.6 s, 3.0 s, and 6.0 s, respectively. The RapidFire 365 system was coupled to an Ion Mobility QTOF MS using an Agilent jet stream orthogonal electrospray ionization source maintained at the following parameters: nitrogen sheath gas, sheath gas temperature, drying gas, drying gas temperature, and nozzle voltage of at 8 L/min, 275°C, 3 L/min, 325°C, and 2 kV respectively. The IM-MS inlet capillary operated at 4 kV, the high-pressure funnel operated at 4.4 Torr with RF at 100 V DC, trapping funnel at 3.8 Torr and 100V DC, and rear funnel at 3.95 Torr and 150 V DC. The IM was pressurized with ultrahigh purity nitrogen, and the drift potential was 1450 V. All data were acquired in positive and negative electrospray mode with a mass range of m/z 50-1700. The Agilent ESI-L low concentration tuning mix solution (G1969-85000, Agilent Technologies) was analyzed daily by direct infusion for single-field CCS calibration (Kurulugama *et al*., 2015).

#### IMS data pre-processing and feature finding

The PNNL-PreProcessor v2020.07.24 (https://omics.pnl.gov/software/pnnl-preprocessor) was used to generate new raw MS files (Agilent MassHunter “.d”) for each sample run with all frames (ion mobility separations) summed into a single frame and apply 3-points smoothing in the ion mobility dimension and noise filtering with a minimum intensity threshold of 20 counts. Single-frame files were converted to mzML using ProteoWizard v3.0.19228 64-bit (Kessner *et al*., 2008). A custom R script was used to set arrival times as a substitute for retention times and generate “LC-MS-like” mzML files. Feature detection was performed in batch mode using MZmine 2 v2.41.2 (Pluskal *et al*., 2010) with the steps: mzML raw data import, mass detector “Wavelet transform” (noise level 20.0, scale level 7, and wavelet window size 0.25), ADAP chromatogram builder (min group size 3, group intensity threshold 50, min highest intensity 100, *m/z* tolerance 0.008 absolute and 10 ppm), chromatogram deconvolution “Local minimum search” (chromatographic threshold 0.02, search min in RT range 0.4, min relative height 0.15, min absolute height 200, peak duration min 0.4 and max 20), isotope grouper (monotonic shape true, max charge 2, representative isotope lowest *m/z*) and CSV data export.

#### CCS calculation and metabolite annotation

Arrival times of detected IMS-MS features were converted to CCS using the autoCCS Python package (https://github.com/PNNL-Comp-Mass-Spec/AutoCCS) which applies the Agilent’s single-field CCS method. CCS values on each run were calibrated using the closest tuning mix infusion run as a reference and the corresponding known CCS values as reported by the Agilent IM-MS Browser v.10.0. IMS-MS features were matched against the experimental CCS-Compendium database (Picache *et al*., 2019) based on tolerances of 10 ppm m/z and 1% CCS. Plant-related compounds in the PlantCyc v15.1.0 (Schläpfer *et al*., 2017) were considered.

